# Arnold tongue entrainment reveals dynamical principles of the embryonic segmentation clock

**DOI:** 10.1101/2021.10.20.465101

**Authors:** Paul Gerald Layague Sanchez, Victoria Mochulska, Christian Mauffette Denis, Gregor Mönke, Takehito Tomita, Nobuko Tsuchida-Straeten, Yvonne Petersen, Katharina F. Sonnen, Paul François, Alexander Aulehla

## Abstract

Living systems exhibit an unmatched complexity, due to countless, entangled interactions across scales. Here we aim to understand a complex system, i.e. segmentation timing in mouse embryos, without a reference to these detailed interactions. To this end, we develop a coarse-grained approach, in which theory guides the experimental identification of the segmentation clock entrainment responses.

We demonstrate period- and phase-locking of the segmentation clock across a wide range of entrainment parameters, including higher-order coupling. These quantifications allow to derive the phase response curve (PRC) and Arnold tongues of the segmentation clock, revealing its essential dynamical properties. Our results indicate that the somite segmentation clock has characteristics reminiscent of a highly non-linear oscillator close to an infinite period bifurcation and suggests the presence of long-term feedbacks.

Combined, this coarse-grained theoretical-experimental approach reveals how we can derive simple, essential features of a highly complex dynamical system, providing precise experimental control over the pace and rhythm of the somite segmentation clock.

## Introduction

How do we gain insight into a complex system, which exhibits emergent properties that reflect the integration of entangled interactions and feedback regulation? As pointed out in the late 1970s by David Marr and Tomaso Poggio in their seminal paper (3), understanding the complexity encountered when studying the “nervous system or a developing embryo” requires the analysis at multiple levels of organization. Their core tenet is that also in biological systems, different levels of organization, while obviously causally linked, exhibit only a loose connection and importantly, can be studied and understood independently from each other.

Such observations are not specific to biology and have been made more quantitative in other fields. In physics, renormalization techniques coarse-grain degrees of freedom to obtain scale-free theories, allowing to define universality classes independent of the precise details of interactions (4, 5). Another recent example is the parameter space compression theory, showing how complex systems (in biology or physics) can be typically reduced to simpler descriptions with few parameters (6–8).

Going one step further, this suggests that one might be able to study- and control-complex systems provided we identify the essential, macro-level behaviour. This is possible because only a limited number of universal descriptions exists, with defining behaviours and properties, that do not depend on the detailed implementation. A central challenge that remains is to implement these theoretical ideas to the experimental study of biological complexity.

Here, we develop a coarse-grained approach combining theory and experiments to study a cellular oscillator ensemble that constitutes the embryonic somite segmentation clock. Functionally, this clock controls the periodic formation of somites, the precursor of the vertebral column and other tissues (9, 10). Molecularly, the segmentation clock comprises the oscillatory activity of several major signaling pathways, such as the Notch, Wnt and Fgf signaling pathways, which show oscillatory dynamics with a period matching somite formation, i.e. ∼ 2 hours in mouse embryos (11–15). More recently, segmentation clock oscillations with a period of ∼ 5 hours have been identified in human induced pluripotent stem cells (iPSCs) differentiated into paraxial mesoderm, identifying a set of ∼ 200 oscillating genes, including targets of Notch and Wnt signaling. (16–18).

Strikingly, as individual oscillating cells are coupled to their neighbours via Notch-Delta signaling, the oscillations occur synchronized and wave-like activity patterns appear to periodically sweep along the embryonic anterior-posterior axis (15, 19–24).

Adding to the complexity, these periodic spatiotemporal wave patterns are linked to an underlying spatial period gradient along the embryonic axis, i.e. signaling dynamics in cells close to the posterior of the embryo oscillate faster compared to those in cells located more anteriorly. Such a period gradient linked to the segmentation clock has been identified in several species (2, 25–30) and also in in vitro assays culturing intact or even dissociated PSM (2, 29).

Of note, an analogous oscillatory system was also described during segmentation in arthropods (31) and while distinct at molecular level, it also exhibits spatiotemporal wave patterns traversing the embryo axis, again with indication of a period gradient (32–35).

In this work, we coarse-grain these underlying complexities and take a dynamical systems, macro-perspective on the segmentation clock, studying it as a single phase-oscillator (Figure 1A). We build on the theory of synchronization and entrainment (see below) to first perform a systematic experimental characterization of its response to perturbation. We compare the outcome to qualitative and quantitative theoretical predictions. In turn, these experimental quantifications allow to derive a phase response curve (PRC) that uniquely characterizes the dynamical properties of the segmentation clock. This new insight provides the means to understand- and control-the timing of a complex embryological patterning process.

**Fig. 1.**
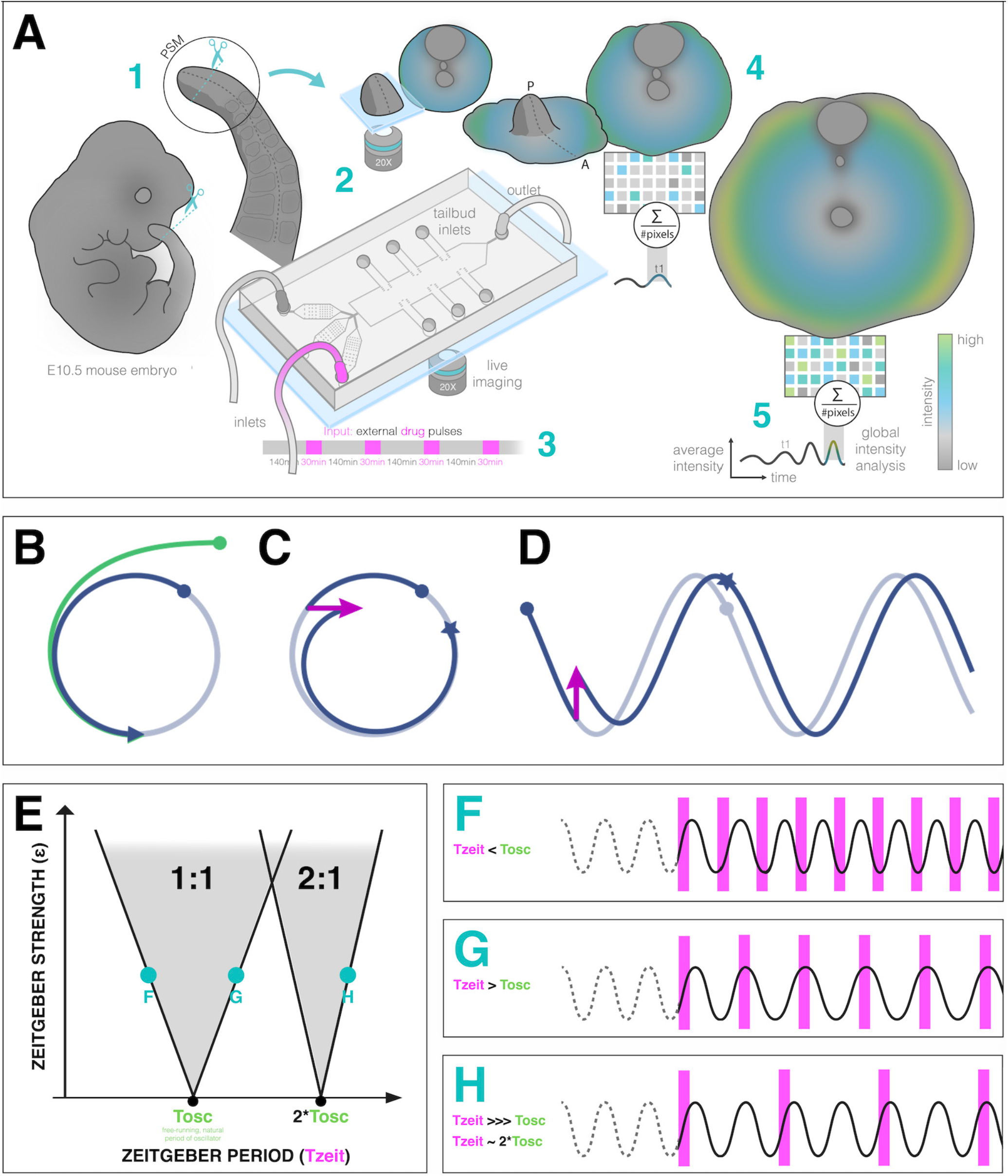
Entrainment of an embryonic oscillator with a *zeitgeber* : setup and theory. (**A**) Schematic of the experimental entrainment setup using a microfluidics device previously described in (1) and an overview of the analysis approach used in this present study. (1) The posterior presomitic mesoderm (PSM) is recovered from E10.5 mouse embryo and used in a quasi-2-dimensional segmentation assay(‘2D-assay’)(2). (2) The embryonic tissue is loaded into a microfluidics device bonded to fibronectin-coated coverglass, and is imaged using a 20x objective at 37^*°*^ C and 5% CO_2_. (3) Simultaneously, the 2D-assay is subjected to periodic pulses of a drug (e.g. DAPT for Notch signaling), which serves as *zeitgeber*. (4) Using dynamic fluorescent reporter for segmentation clock genes (e.g. LuVeLu for Notch signaling), we quantify endogenous signaling oscillations during the experiment. (5) We generate a single time-series using reporter signal from the entire sample (‘global ROI’) that allows to quantify segmentation clock rate and rhythm (see validation Figure 2 and Supplementary Figure S17. Illustration by Stefano Vianello. A photo of the actual microfluidics device and its design are shown in Figure S1. (**B**) Abstract definition of phase: two different points in the plane (*x, y*) have the same phase if they converge on the same point on the limit cycle (indicated in grey). (**C**) Perturbations change the phase of the cycle by an increment Δ *φ* (here phase is defined by the angle in the plane). (**D**) Time courses with similar perturbation as C showing the oscillations of *x* and phase difference Δ *φ*. (**E**) Illustration of generic Arnold tongues, plotted as a function of *zeitgeber* strength (ε) and *zeitgeber* period (*T*_*zeit*_), mapping *n* : *m* entrainment where the entrained oscillator (with natural period of *T*_*osc*_) goes through *n* cycle/s for every *m* cycle/s of the *zeitgeber*. Three different points in the 1 : 1 and 2 : 1 Arnold tongues are specified with corresponding graphical illustration of an autonomous oscillator as it is subjected to *zeitgeber* with different periods (*T*_*zeit*_): when *T*_*zeit*_ is less than *T*_*osc*_ (**F**), when *T*_*zeit*_ is greater than *T*_*osc*_ (**G**), and when *T*_*zeit*_ is much much greater than *T*_*osc*_ but is close to twice of *T*_*osc*_ (**H**). Free-running rhythm of the oscillator (i.e. before perturbation) is marked by a dashed line, while solid line illustrates its rhythm during perturbation with *zeitgeber*. Magenta bars represent the *zeitgeber* pulses. Illustration by Stefano Vianello. A high-resolution version of this figure is available at https://doi.org/10.6084/m9.figshare.19093112.

### Theory of synchronization guides the experimental study of segmentation clock entrainment

Our experimental study is based on and guided by the theory of entrainment of oscillators by external periodic signal -a subset of the general theory of synchronization (36).

Entrainment is observed when an autonomous oscillator adapts its behaviour to lock to an external periodic signal (called *zeitgeber* in the circadian rhythm literature). The general theoretical framework to understand entrainment requires the definition of oscillator phases (Figure 1B-D), and their response to perturbation (37). Assuming the *zeitgeber* consists in periodic pulses, entrainment is observed when the phase of the oscillator *φ*_*ent*_ at the time of the *zeitgeber* is constant (technically, a fixed point of the Poincaré return map (38)). This defines period-locking (also termed mode locking) (36).

Entrainment is not always manifested and conditions for its existence can be derived. Quantitatively, when entrainment occurs, the *zeitgeber* induces a periodic phase perturbation (or response) of the entrained oscillator, which *exactly* compensates the detuning (or period mismatch, *T*_*zeit*_ *− T*_*osc*_) between the *zeitgeber* and the free-running oscillator. For this reason, when the detuning is very small, a weak external perturbation is enough to entrain an oscillator. Conversely, if the detuning is big, a strong signal and associated response is required for entrainment. One can then plot the miminal strength of the *zeitgeber* (ε) versus corresponding detuning (or simply *T*_*zeit*_, if *T*_*osc*_ is constant): these maps are more commonly known as Arnold tongues (Figure 1E). Arnold tongues predict the period- and phase-locking behaviour in oscillatory systems as different as electrical circuits (39), oscillatory chemical reactions (40–43), or living systems like circadian rhythms (44).

Lastly, more complex patterns of entrainment can be observed: for instance, stable phase relationships can be established where the entrained oscillator goes through *n* cycles for every *m* cycle of the external signal, defining *n* : *m* period-locking. In that case the instantaneous period of the oscillator matches *m/n* the period of the *zeitgeber* (*T*_*zeit*_). Corresponding Arnold tongues can be obtained, leading to a rich structure for entrainment in parameter space (Figure 1F-H).

Previously, periodic activation of Notch signaling via heat shock-driven expression of the ligand Delta was used to modulate the segmentation of PSM in zebrafish (45). In the cited study, the readout to assess the effect of the periodic perturbation relied on somite size and morphology. While this gave important insight into how morphological segmentation of the PSM and somite length are affected by a *zeitgeber*, experimental investigation on the underlying signaling dynamics upon periodic perturbation remains limited.

In this work, to experimentally apply the theory of synchronization to the segmentation clock, we make use of a microfluidics-based entrainment setup, which we had established previously in the lab (Figure S1) (1).

We showed before that using a quasi-2-dimensional in vitro segmentation assay (hereafter referred to as a 2D-assay), which recapitulate segmentation clock dynamics and PSM patterning (2), the microfluidics-entrainment approach allowed us to take control of Notch and Wnt signaling oscillations, providing direct functional evidence that the oscillation phase shift between Wnt and Notch signaling is critical for PSM patterning (1).

## Results

### A coarse-grained, single oscillator description of the segmentation clock

To perform a systematic analysis of entrainment dynamics, we first introduced a single oscillator description of the segmentation clock. We used the segmentation clock reporter LuVeLu, which shows highest signal levels in regions where segments form (15). Hence, we reasoned that a global ROI quantification, averaging LuVeLu intensities over the entire sample, should faithfully report on the segmentation rate and rhythm, essentially quantifying ‘wave arrival’ and segment formation in the periphery of the sample. Indeed, our validation confirmed that the oscillation period based on the global ROI analysis using the LuVeLu reporter closely matched the rate of morphological segment boundary formation(Figure 2D, S2). In addition, global ROI quantifications using additional reporters to measure Wnt signalling oscillations (i.e. Axin2, Figure 2C-D, S2) and the segmentation marker Mesp2 (Figure 2D, S2) also showed close correspondence with LuVeLu global ROI measurements of segmentation clock period.

**Fig. 2.**
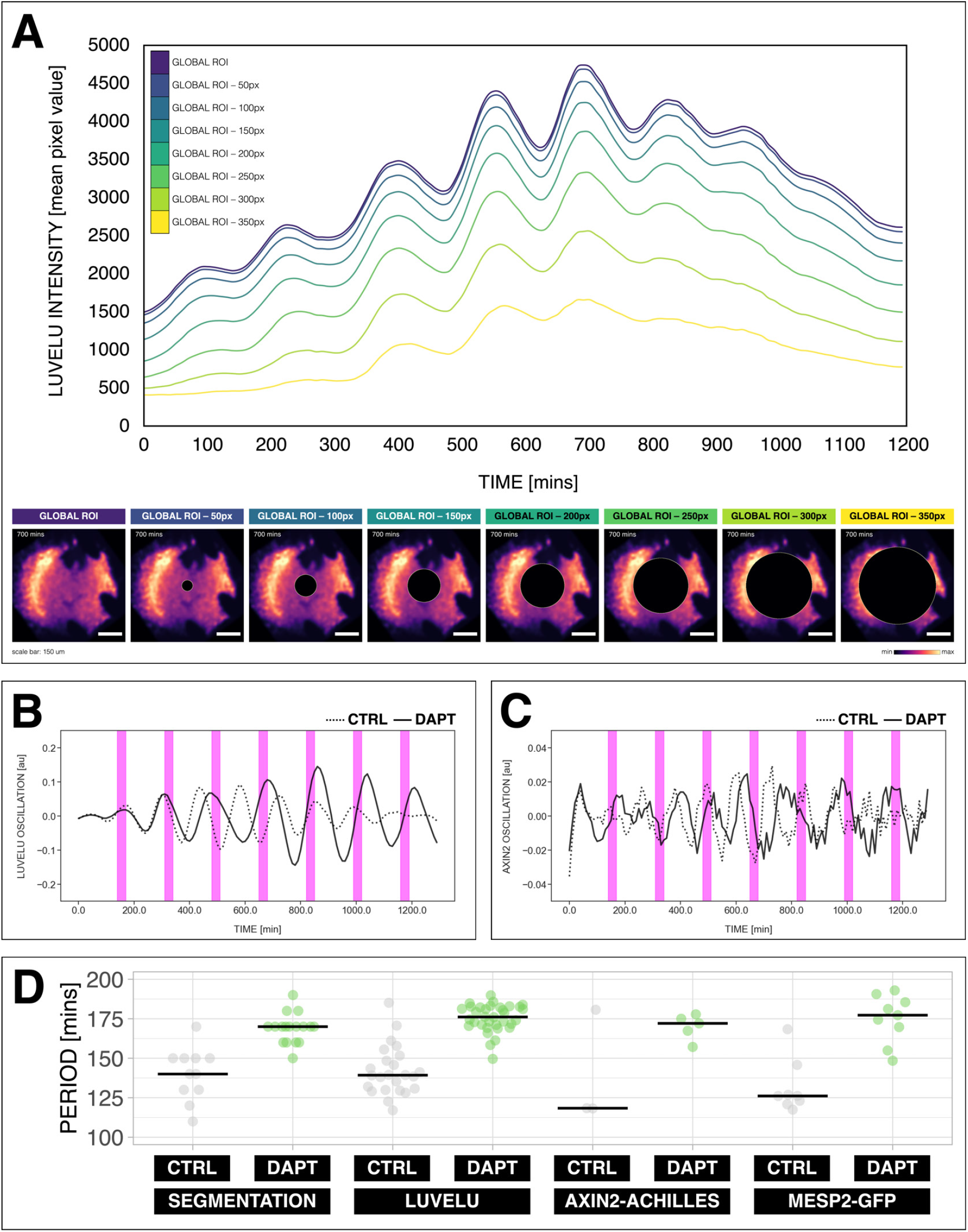
Quantifying the segmentation clock rhythm. (**A**) Comparison of measurements obtained from a region of interest (ROI) spanning the entire field (“global ROI”) with those obtained from ROIs in which central regions of increasing size were excluded. The excluded area is represented in terms of the diameter (in pixels, px) of a circular region at the center. The corresponding timeseries are shown in the top panel and marked with different colors. Bottom panel shows the global ROI and excluded region in a snapshot of signal of LuVeLu, a dynamic reporter of Notch signaling driven from the Lfng promoter (15), in a 2D-assay at 700 mins from the start of the experiment. Henceforth, timeseries is obtained using “global ROI”, unless otherwise specified. (**B**) Detrended timeseries of the segmentation clock (obtained using global ROI) in 2D-assays, which express the LuVeLu reporter, subjected to 170-min periodic pulses (magenta bars) of either 2 uM DAPT (in solid line) or DMSO (for control, in dashed line). Timelapse movie and corresponding timeseries from global ROI are available at https://youtu.be/fRHsHYU_H2Q. (**C**) Detrended timeseries of Axin2-linker-Achilles reporter in 2D-assays, subjected to 170-min periodic pulses (magenta bars) of either 2 uM DAPT (in solid line) or DMSO (for control, in dashed line). (**D**) Period of morphological segment boundary formation, LuVeLu oscillation, Axin2-linker-Achilles oscillation, and rhythm of Mesp2-GFP in 2D-assays subjected to 170-min periodic pulses of either 2 uM DAPT (or DMSO for control). Each sample is represented as a dot, the median is denoted as a solid horizontal line. For morphological segment boundary formation, period was determined by taking the time difference between two consecutive segmentation events (for CTRL: 17 segmentation events in 6 samples from 4 independent experiments, and for DAPT: 24 segmentation events in 7 samples from 5 independent experiments, with p-value < 0.001) in the brightfield channel. For period quantifications based the reporters, the mean period per sample from 650 to 850 mins after start of the experiment was plotted. LuVeLu: (CTRL: n = 24 and N = 7) and (DAPT: n = 34 and N = 8) with p-value < 0.001, Axin2-linker-Achilles: (CTRL: n = 3 and N = 1) and (DAPT: n = 5 and N = 1) with p-value = 0.107, Mesp2-GFP: (CTRL: n = 8 and N = 3) and (DAPT: n = 9 and N = 3) with p-value = 0.01. Data were visualized using PlotsOfData (46). To calculate the p-value, two-tailed test for absolute difference between medians was done via a randomization method using PlotsOfDifferences (47). The timeseries and corresponding period evolution during entrainment, obtained from wavelet analysis, are in Figure S2A and Figure S2B, respectively. Timelapse movies are available at https://youtu.be/edFczx_-9hM and https://youtu.be/tQeBk0_U_Qo, respectively. A high-resolution version of this figure is available at https://doi.org/10.6084/m9.figshare.19093112.

Hence, we conclude that LuVeLu global ROI timeseries analysis provides a valid coarse-grained quantification of the pace (period ∼ rate of segment formation) and rhythm (phase) of the segmentation clock.

### The pace of the segmentation clock can be locked to a wide range of entrainment periods

Having established a quantitative, coarse-grained read-out for segmentation clock pace and rhythm, we next analyzed whether pace and rhythm can be experimentally tuned using microfluidics-based entrainment (Figure 1A, S1).

First, we tested whether the segmentation clock can be entrained to periods different and far from the endogenous period of ∼ 140 mins, which we refer to as its free-running, natural period or *T*_*osc*_). To address this question, we modified the entrainment period from 120 to 180 minutes, while keeping the drug concentration and the pulse duration (i.e. 30 mins/cycle) constant. Our results show that, while controls cycled close to *T*_*osc*_ (Figure 3A-C, S3), the segmentation clock rhythm in DAPT-entrained samples closely adjusted to *T*_*zeit*_ over the specified range (Figure 3D, S3-S4). Hence, we were able to speed up and slow down the pace of the segmentation clock system using entrainment.

**Fig. 3.**
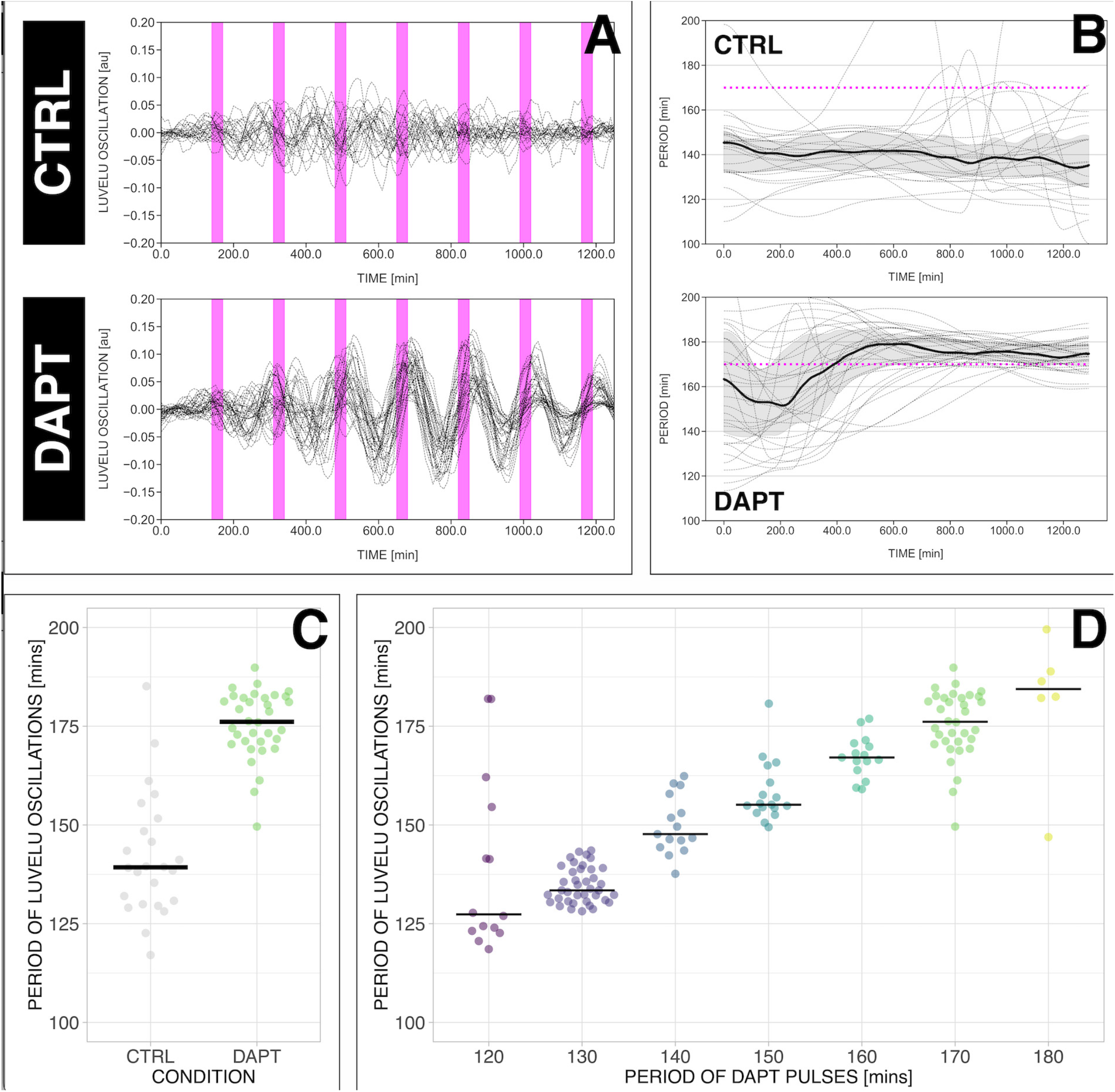
The segmentation clock can be locked to a wide range of entrainment periods. (**A**) Detrended (via sinc-filter detrending) timeseries of the segmentation clock in 2D-assays subjected to 170-min periodic pulses of 2 uM DAPT (or DMSO for controls). Periodic pulses are indicated as magenta bars and the timeseries of each sample (for CTRL: n = 24 and N = 7, for DAPT: n = 34 and N = 8) is marked with a dashed line. (**B**) Period evolution during entrainment, obtained from wavelet analysis. The period evolution for each sample and the median of the periods are represented here as a dashed line and a solid line, respectively. The gray shaded area corresponds to the interquartile range. Magenta dashed line marks *T*_*zeit*_. (**C**) Mean period from 650 to 850 mins after start of the experiment of samples subjected to 170-min periodic pulses of 2 uM DAPT (or DMSO for controls), with p-value < 0.001. Each sample is represented as a dot, while the median of all samples is denoted as a solid horizontal line. This plot is the same as the plot for the LuVeLu condition in Figure 2D. (**D**) Mean period from 650 to 850 mins after start of the experiment of samples entrained to periodic pulses of 2 uM DAPT. Each sample is represented as a dot, while the median of all samples is denoted as a solid horizontal line. The period of the DAPT pulses is specified (for 120-min: n = 14 and N = 3, for 130-min: n = 39 and N = 10, for 140-min: n = 15 and N = 3, for 150-min: n = 17 and N = 4, for 160-min: n = 15 and N = 3, for 170-min: n = 34 and N = 8, for 180-min: n = 6 and N = 1). Data were visualized using PlotsOfData (46), and a summary is provided in Table S1. A similar plot including each condition’s respective control is in Figure S3. The analysis of period and wavelet power across time is summarized in Figure S4. To calculate the p-value, two-tailed test for absolute difference between medians was done via a randomization method using PlotsOfDifferences (47). A high-resolution version of this figure is available at https://doi.org/10.6084/m9.figshare.19093112.

Notably, period-locking was less precise (i.e. higher standard deviation as shown in Table S1) with 120-min and 180-min *zeitgeber* periods, a possible indication that the limit of entrainment range is approached at these conditions.

We also tested the effect of varying DAPT concentration, as means of changing *zeitgeber* strength (ε), on entrainment dynamics. Synchronization theory predicts that *zeitgeber* strength correlates with the time it takes to reach period-locking (36, 48). To test this prediction, we entrained samples with periodic DAPT pulses at fixed intervals of 170 minutes and varied drug concentration. We indeed found that the time needed to show period-locking was shortened in samples using higher DAPT concentrations (Figure 4A-B, S5A). Additionally, as expected, higher drug concentrations also resulted in more robust entrainment, indicated by the quantifications of the first Kuramoto order parameter *R*, a measure for in-phase synchrony between samples (Figure 4C-D, 3A, S5B) (see definition in Supplement, notice that *R* is the modulus of the mean-field variable used in (49)).

**Fig. 4.**
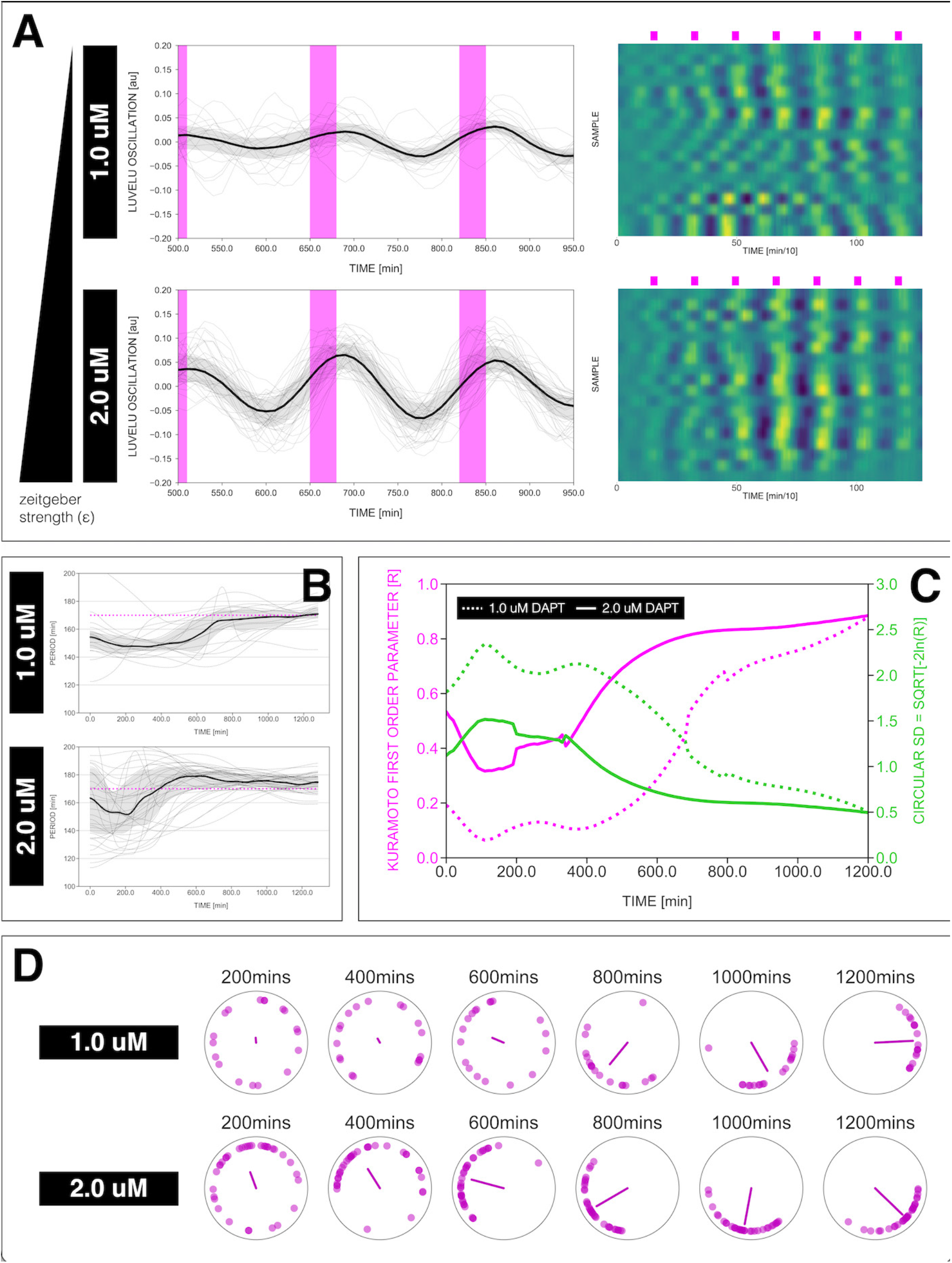
Effect of varying DAPT concentrations on entrainment dynamics. (**A**) Left: Detrended (via sinc-filter detrending) timeseries of the segmentation clock in 2D-assays entrained to 170-min periodic pulses of either 1 uM or 2 uM DAPT, zoomed in from 500 mins to 950 mins. Periodic pulses are indicated as magenta bars and the timeseries of each sample (for 1 uM: n = 18 and N = 5, for 2 uM: n = 34 and N = 8) is marked with a dashed line. The median of the oscillations is represented here as a solid line, while the gray shaded area denotes the interquartile range. Right: Detrended (via sinc-filter detrending) timeseries of the segmentation clock in 2D-assays entrained to 170-min periodic pulses of either 1 uM (n = 18 and N = 5) or 2 uM (n = 18 and N = 8) DAPT represented as heatmaps, generated using PlotTwist (50). Periodic pulses are indicated as magenta bars. Each row corresponds to a sample. (**B**) Period evolution during entrainment, obtained from wavelet analysis. The period evolution for each sample and the median of the periods are represented here as a dashed line and a solid line, respectively. The gray shaded area corresponds to the interquartile range. The plot for the 2 uM condition is the same as the plot for the DAPT condition in Figure 3B. Magenta dashed line marks *T*_*zeit*_. (**C**) Evolution of first Kuramoto order parameter (*R*) in magenta and circular standard deviation (*circSD*) in green over time, showing change in coherence of multiple samples during the experiment. An *R* equal to 1.0 means that samples are in-phase. *circSD* is equal to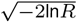. (**D**) Polar plots at different timepoints showing phase of each sample and their first Kuramoto order parameter, represented as a magenta dot along the circumference of a circle and a magenta line segment at the circle’s center, respectively. A longer line segment corresponds to a higher first Kuramoto order parameter, and thus to more coherent samples. The direction of the line denotes the vectorial average of the sample phases. Time is indicated as mins elapsed from the start of the experiment. A high-resolution version of this figure is available at https://doi.org/10.6084/m9.figshare.19093112.

### Higher-order entrainment of the segmentation clock

In theory, a non-linear oscillator should be amenable not only to 1 : 1 entrainment, but also to higher-order *n* : *m* entrainment, in which *n* cycles of the endogenous oscillation lock to *m* cycles of the *zeitgeber* (36, 51). In practice, demonstration of higher-order entrainment is challenging due to the narrow permissive parameter region, i.e. Arnold tongues are progressively narrower away from the 1 : 1 regime. Strikingly, we found experimental evidence for higher-order 2 : 1 entrainment (Figure 5). Samples entrained with either 300-min or 350-min pulses showed evidence of 2 : 1 entrainment (Figure 5A-C), i.e. the segmentation clock oscillated twice per each *zeitgeber* pulse and hence the segmentation clock rhythm adjusted to a period very close to 150 mins for *T*_*zeit*_ = 300 mins and approaching 175 mins for *T*_*zeit*_ = 350 mins (Figure 5D). We also found evidence for phase-locking during 2 : 1 entrainment, i.e. a narrowing of the phase distribution amongst the individual samples ((Figure 5C).

**Fig. 5.**
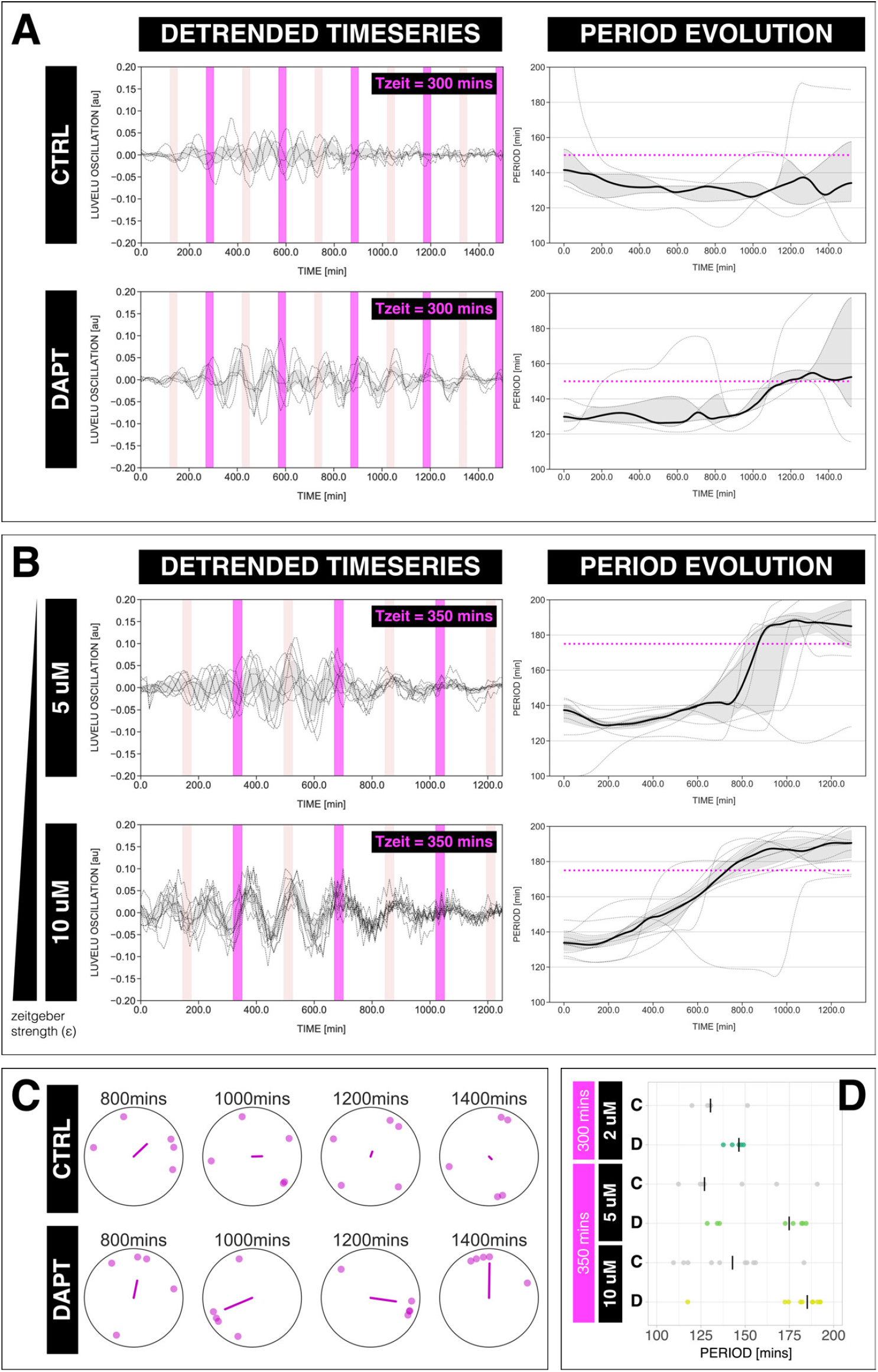
The segmentation clock can be entrained to a higher order. (**A**) Left: Detrended (via sinc-filter detrending) timeseries of the segmentation clock in 2D-assays subjected to 300-min periodic pulses of 2 uM DAPT (or DMSO for controls). Periodic pulses are indicated as magenta bars and the timeseries of each sample (for CTRL: n = 5 and N = 1, for DAPT: n = 5 and N = 1) is marked with a dashed line. The gray shaded area denotes the interquartile range. Hypothetical pulses at half the *zeitgeber* period are indicated as light pink bars. Right: Period evolution during entrainment, obtained from wavelet analysis. The period evolution for each sample and the median of the periods are represented here as a dashed line and a solid line, respectively. The gray shaded area corresponds to the interquartile range. Magenta dashed line marks 0.5*T*_*zeit*_. (**B**) Left: Detrended (via sinc-filter detrending) timeseries of the segmentation clock in 2D-assays subjected to 350-min periodic pulses of either 5 uM DAPT or 10 uM DAPT. Periodic pulses are indicated as magenta bars and the timeseries of each sample (for 5 uM DAPT: n = 8 and N = 2, for 10 uM DAPT: n = 10 and N = 2) is marked with a dashed line. The gray shaded area denotes the interquartile range. Hypothetical pulses at half the *zeitgeber* period are indicated as light pink bars. Right: Period evolution during entrainment, obtained from wavelet analysis. The period evolution for each sample and the median of the periods are represented here as a dashed line and a solid line, respectively. The gray shaded area corresponds to the interquartile range. Magenta dashed line marks 0.5*T*_*zeit*_. (**C**) Polar plots at different timepoints showing the phase of each sample in (A) and the first Kuramoto order parameter, represented as a magenta dot along the circumference of a circle and a magenta line segment at the circle’s center, respectively. A longer line segment corresponds to a higher first Kuramoto order parameter, and thus to more coherent samples. The direction of the line denotes the vectorial average of the sample phases. Time is indicated as mins elapsed from the start of the experiment. (**D**) Mean period either from 1000 to 1150 mins (for *T*_*zeit*_ = 300 mins) or from 800 to 950 mins (for *T*_*zeit*_ = 350 mins) after start of the experiment of samples subjected to periodic pulses of DAPT (or DMSO for controls). Each sample is represented as a dot, while the median of all samples is denoted as a solid vertical line. For 300-min 2 uM DAPT: (CTRL: n = 5 and N = 1) and (DAPT: n = 5 and N = 1) with p-value = 0.191, for 350-min 5 uM DAPT: (CTRL: n = 7 and N = 2) and (DAPT: n = 8 and N = 2) with p-value = 0.049, for 350-min 10 uM DAPT: (CTRL: n = 10 and N = 2) and (DAPT: n = 10 and N = 2) with p-value = 0.016. The period and concentration of the DAPT pulses are specified. Data were visualized using PlotsOfData (46). To calculate p-values, two-tailed test for absolute difference between medians was done via a randomization method using PlotsOfDifferences (47). A high-resolution version of this figure is available at https://doi.org/10.6084/m9.figshare.19093112.

### The segmentation clock establishes a stable entrainment phase relative to the *zeitgeber*

We next analyzed the phase-locking behaviour between the segmentation clock and the *zeitgeber*. According to dynamical systems theory, phase-locking, i.e. the entrainment phase (*φ*_*ent*_), can be characterized as an attractor, i.e. it is a stable fixed point (38, 52). To quantify the entrainment phase, we plotted the data as stroboscopic maps (Figure 6A), which take a snapshot of the to the posterior PSM), the entrainment phase at *T*_*zeit*_ = 130 segmentation clock phase at regular intervals based on *zeitgeber* pulses (38, 52, 53). Stroboscopic maps enable determination of *φ*_*ent*_ as a stable fixed point that lies on the diagonal, where there is phase-locking.

**Fig. 6.**
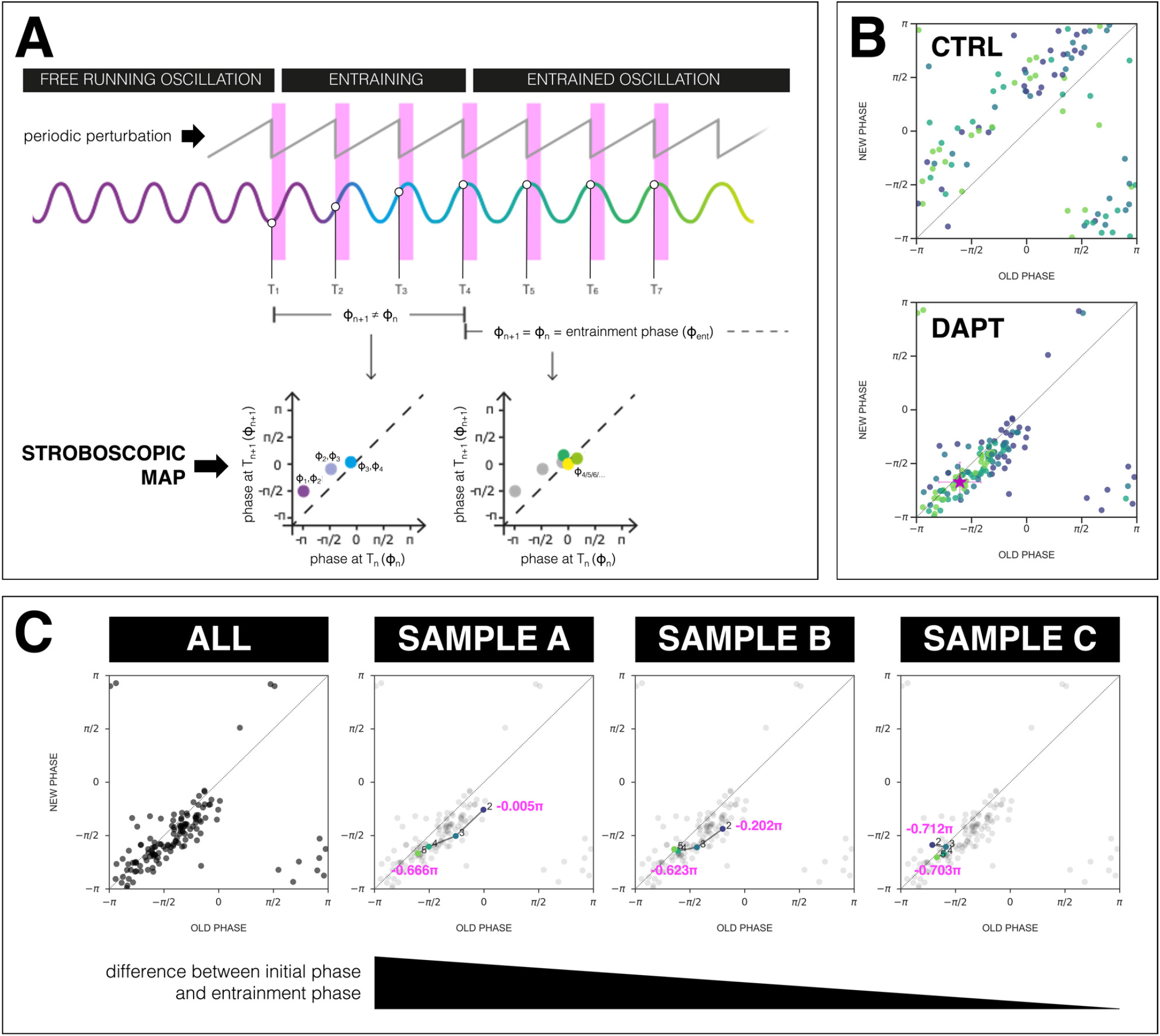
The segmentation clock establishes a stable phase relationship with the *zeitgeber*. (**A**) Schematic of how to generate a stroboscopic map, where the phase of the segmentation clock just before a DAPT pulse (old phase, *φ*_*n*_) is iteratively plotted against its phase just before the next pulse (new phase, *φ*_*n*+1_).The position of each point in a stroboscopic map thus denotes a stepwise change in phase of the segmentation clock as it undergoes entrainment to the *zeitgeber*. Upon entrainment and phase-locking, the new phase is equal to the old phase lies on the diagonal of the stroboscopic map. This point on the diagonal is the entrainment phase (*φ*_*ent*_). Illustration by Stefano Vianello. (**B**) Stroboscopic map of samples subjected to 170-min periodic pulses of 2 uM DAPT (or DMSO for controls). Colors mark progression in time, from purple (early) to yellow (late). Note that while in control samples, points remain above the diagonal (reflecting that endogenous oscillations run faster than *T*_*zeit*_ = 170 mins as shown in Figure 3B-C), in entrained samples, the measurements converge towards a point on the diagonal (i.e. the entrainment phase, *φ*_*ent*_), showing phase-locking. This localized region marks the entrainment phase (*φ*_*ent*_). This is highlighted with a magenta star, which corresponds to the centroid (*x*_*c*_, *y*_*c*_). The centroid was calculated from the vectorial average of the phases of the samples at the end of the experiment, where *x*_*c*_ = vectorial average of old phase, *y*_*c*_ = vectorial average of new phase. The spread of the points in the region is reported in terms of the circular standard deviation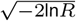, where *R* is the first Kuramoto order parameter). (**C**) Stroboscopic maps of the segmentation clock entrained to 170-min periodic pulses of 2 uM DAPT (n = 34 and N = 8), for all samples (ALL) and for three individual samples (SAMPLE A, SAMPLE B, SAMPLE C). The numbers and colors (from purple to yellow) denote progression in time. The initial phase (old phase at point 2) and the entrainment phase (new phase at point 5) of each sample are specified. A high-resolution version of this figure is available at https://doi.org/10.6084/m9.figshare.19093112.

Plotting the stroboscopic maps shows that in samples entrained to periodic pulses of DAPT the segmentation clock phase dynamics converge to a region on the diagonal, which marks *φ*_*ent*_ (Figure 6B). Such convergence towards a stable entrainment phase is further highlighted by looking at consecutive pulses and the corresponding phase trajectories of individual samples. As exemplified in Figure 6C and as previously theoretically predicted (48, 54), at the same *zeitgeber* strength and *zeitgeber* period, faster (or slower) convergence towards this fixed point (i.e. entrainment) was achieved when the initial phase of the endogenous oscillation (*φ*_*init*_) was closer (or farther) to *φ*_*ent*_.

### The phase of entrainment varies according to *zeitgeber* period

One fundamental property of entrained oscillatory systems is that its phase of entrainment varies as a function of the detuning, i.e. the period mismatch between endogenous oscillator and entrainment period (44, 55–58). In addition, theoretical studies have supported a “180-degree” rule (44), stating that the phase of entrainment varies by half a cycle (180° or π) within the range of permissible *zeitgeber* entrainment periods, *T*_*zeit*_ (44, 56).

To test these theoretical predictions, we quantified *φ*_*ent*_ across a wide range of detuning, i.e. from 120-min to 180-min *zeitgeber* period. We found that *φ*_*ent*_, based on the centroid localization close to the diagonal in the stroboscopic maps, gradually changed its position as *zeitgeber* period, i.e. detuning, was varied (Figure 7A, S9). From 120-min to 180-min entrainment periods, *φ*_*ent*_ systematically shifted from ∼ π/2 to ∼ 3π/2 (Figure 7A-B), spanning a range of almost π(Figure 7B-C). Hence, as predicted by theory, our results show that the segmentation clock phase of entrainment varies as a function of the detuning relative to zeitgeber pulses. It was previously documented that the period of tissue seg_-mentation matches the period of oscillations localized in the posterior PSM, i.e. center of of the 2D-assays (29, 45). We were then curious whether or not the dynamics from our coarse-grained analysis match that for oscillations in the center of the 2D-assays, and if the behaviour of oscillations in this localized region also exhibits dependence to *zeitgeber* _ period. To study this, we compared the timeseries acquired from (a) global ROI spanning the entire field of view and from (b) a circular ROI restricted at the center of the 2D_ assays (“center ROI”, radius: 25 pixels, 1 pixel = 1.38 um). Upon comparison, we noted that the oscillations at the center of 2D-assays adjusted its rhythm to the zeitgeber with strong correspondence to the coarse-grained oscillator (Figure S6), regardless whether the segmentation clock was sped up (Figure S6A-B) or slowed down (Figure S6C-D). Moreover, the phase-locking between the zeitgeber and either the segmentation clock or the oscillations at the center of the samples are similar (Figure S6B1-B2, D1-D2, S7). Concomitantly, to the posterior PSM), the entrainment phase at T_*zeit*_ = 130mins is different to the entrainment phase at *T*_*zeit*_ = 170 mins (Figure S7B), here too exemplifying the effect of *zeitgeber* period on the phase of entrainment.

**Fig. 7.**
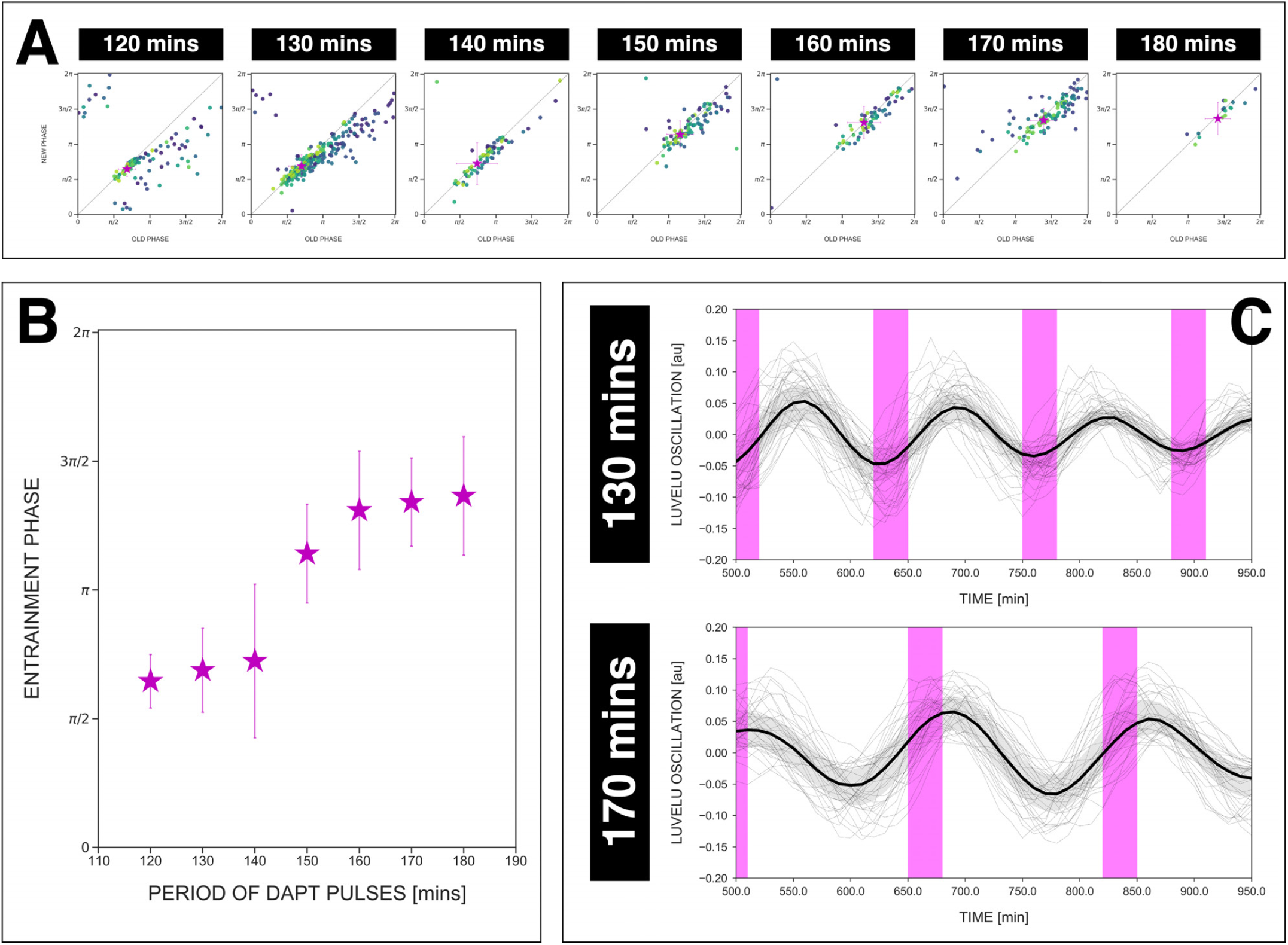
The entrainment phase varies according to *zeitgeber* period within a range of π. (**A**) Stroboscopic maps for different periods of DAPT pulses (i.e. *zeitgeber* periods) placed next to each other. In these maps, only samples that were phase-locked by the end of the experiment are considered (for 120-min: n = 13/14 and N = 3/3, for 130-min: n = 38/39 and N = 10/10, for 140-min: n = 10/15 and N = 3/3, for 150-min: n = 16/17 and N = 4/4, for 160-min: n = 11/15 and N = 3/3, for 170-min: n = 28/34 and N = 8/8, for 180-min: n = 4/6 and N = 1/1). Here, a sample was considered phase-locked if the difference between its phase at the time of the final drug pulse considered and its phase one drug pulse before is less than *π/*8. The localized region close to the diagonal in each map marks the entrainment phase (*φ*_*ent*_) for that *zeitgeber* period. This is highlighted with a magenta star, which corresponds to the centroid (*x*_*c*_, *y*_*c*_). The centroid was calculated from the vectorial average of the phases of the samples at the end of the experiment, where *x*_*c*_ = vectorial average of old phase, *y*_*c*_ = vectorial average of new phase. The spread of the points in the region is reported in terms of the circular standard deviation 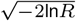, where *R* is the first Kuramoto order parameter). The *zeitgeber* period is indicated above the maps. Colors mark progression in time, from purple to yellow. Stroboscopic maps of all samples and their respective controls are shown in Figure S9. Drug concentration and drug pulse duration were kept constant between experiments at 2 uM and 30 mins/cycle, respectively. Phase 0 is defined as the peak of the oscillation. (**B**) Entrainment phase (*φ*_*ent*_) at different *zeitgeber* periods, each calculated from the vectorial average of the phases of phase-locked samples at the time corresponding to last considered DAPT pulse. The spread of *φ*_*ent*_ between samples is reported in terms of the circular standard deviation 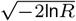, where *R* is the first Kuramoto order parameter). (**C**) Detrended (via sinc-filter detrending) timeseries of the segmentation clock in 2D-assays entrained to either 130-min or 170-min periodic pulses of 2 uM DAPT, zoomed in from 500 mins to 950 mins. Periodic pulses are indicated as magenta bars and the timeseries of each sample (for 130-min: n = 39 and N = 10, for 170-min: n = 34 and N = 8) is marked with a dashed line. The median of the oscillations is represented here as a solid line, while the gray shaded area denotes the interquartile range. The full detrended timeseries for the 170-min condition can be seen in Figure 3A. A high-resolution version of this figure is available at https://doi.org/10.6084/m9.figshare.19093112.

### Phase response curves derivation and period change

Building on our finding that the phase of entrainment varies as a function of detuning (Figure 7B), our goal was to ex-tract the quantitative information embedded in this dynamic behaviour. This is possible since the phase of entrainment dynamics reflect, at quantitative level, the fundamental properties of a dynamical system that responds to external perturbations. This behaviour, in turn, can usually be captured with a single function, the phase response curve (PRC) (44, 52). The PRC describes the change of phase induced by a perturbation, and a priori depends on both the nature of the perturbation received and the phase of the cycle. Because of the direct dependence of the phase of entrainment dynamics on the PRC, our goal was to use the experimental data to gain insight into the segmentation clock PRC.

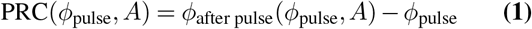

where *φ*_pulse_ is the phase of the segmentation clock *on the cycle* at the moment of the pulse, and *φ*_after pulse_ the phase after the pulse (which can be defined even if the system transiently moves outside of the limit cycle, via isochrons, see Winfree (59)). If, following a perturbation, the oscillator relaxes quickly towards the limit cycle, one can use the PRC to compute response to periodic pulses with period *T*_*zeit*_. The sequence of phases at each pulse is then given by the stroboscopic map, introduced in Figure 6A:

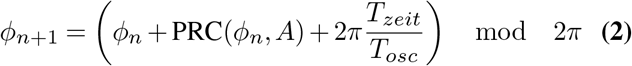

and 1 : 1 entrainment occurs when this stroboscopic map converges towards a single fixed point *φ*_*ent*_ for a given *T*_*zeit*_. When *T*_*zeit*_ is varied, different phases of entrainment *φ*_*ent*_(*T*_*zeit*_) are observed, here plotted in (Fig.7B).

To derive the PRC directly from the experimental stroboscopic maps, we invert Eq. 2 into :

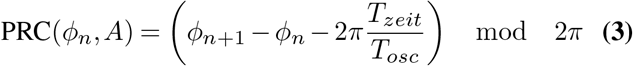

By estimating *φ*_*n*_, *φ*_*n*+1_ for a given *T*_*zeit*_, one can estimate the PRC. The advantage of this approach is that it allows to estimate PRC at any observed phase at the pulse *φ*_*n*_, even far from the entrainment phase *φ*_*ent*_. Notice, however, that phases far from *φ*_*ent*_ will be only sampled over the first few pulses (so with possibly much noise) while conversely, *φ*_*ent*_ will be quickly oversampled (and as such better defined statistically).

Figure 8A1 shows the PRC computed from the data as well as Fourier series fits for different entrainment periods. PRCs computed for different entrainment periods have a similar shape, with similar locations for maxima and minima. Strikingly, those PRCs are not sinusoidal but essentially 0 or strongly negative, an unusual situation from a dynamical systems theory standpoint associated to special classes of oscillators (see more details below and in Discussion). However, contrary to theoretical predictions, the inferred PRCs at different entrainment periods do not overlap and appear shifted vertically as *T*_*zeit*_ is changed.

**Fig. 8.**
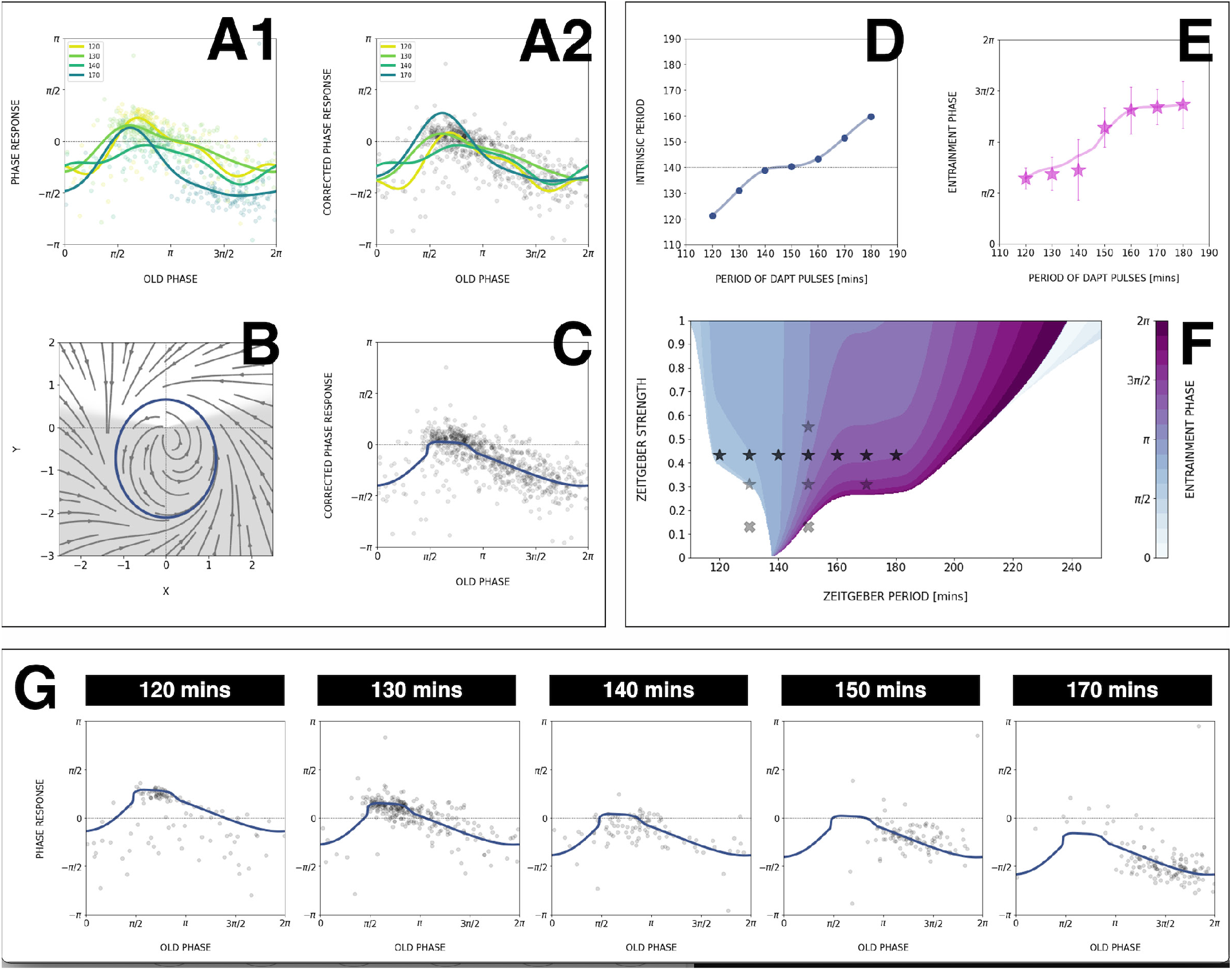
Modelling segmentation clock entrainment response. (**A**) PRC from the data for different *zeitgeber* periods. (**1**) PRCs calculated at different *T*_*zeit*_ (points) and Fourier series fitted to them (lines). (**2**) Original PRCs are shifted vertically to collapse the data points on one curve. (**B**) Oscillator model, optimized by fitting the vertically shifted PRC. The limit cycle is an ellipse (blue) with eccentricity *λ* = 0.5, the region with speeding up *s*_*\**_ = 5.6 is shaded. (**C**) Optimized model PRC (line) overlaid to the vertically shifted data points. (**D**) Modelled intrinsic period *T*_*osc*_ as a function of entrainment period *T*_*zeit*_. Points were inferred from the PRCs data in **A1** by matching the detuning and entrainment phase *φ*_*ent*_, the function was interpolated with cubic splines. (**E**) Entrainment phase *φ*_*ent*_ from the model with changing intrinsic period (line) agrees with the experimental data (points and error bars). (**F**) 1 : 1 Arnold tongue and isophases calculated with the model. Stars correspond to experimental data (*T*_*zeit*_, *A*) with observed entrainment, in agreement with *φ*_*ent*_ data for different DAPT concentrations (Figure S10A). Black stars represent the experiments used for optimizing the model. Crosses correspond to experimental conditions with no entrainment. (**G**) Fit of the model (solid lines) overlayed with experimental phase differences (dots) at different periods *T*_*zeit*_. A high-resolution version of this figure is available at https://doi.org/10.6084/m9.figshare.19093112.

Such vertical shifts in PRCs as a function of entrainment period have been previously observed in so-called “overdrive suppression” for cardiac cell oscillators (60). Shifts occur when the entrainment signal impacts the biochemical control of the system, leading to a change of intrinsic period (here *T*_*osc*_). Here, such change of intrinsic period can not be directly measured experimentally, as the system is entrained to another period, however, we found several lines of experimental and theoretical evidence, in addition to the vertical PRC shift, in full agreement with such an effect.

First, the slope of the *φ*_*ent*_(*T*_*zeit*_) curve in Figure 7B is unusual: PRC theory predicts that for high and low detuning this curve should have vertical slopes. Here, we observed plateaus in the data, which can be simply explained if the intrinsic period changes (see mathematical explanation in Supplementary Note 2). In addition, we performed entrainment release experiments, where the segmentation clock is first entrained, then released (Figure S8): we observed a slow recovery over several cycles to a period matching control samples, compatible with a transient change of the intrinsic period. This effect was confirmed by a more precise study of the phase return map after release (Figure S8B). Combined, the inference of the PRC based on entrainment quantification data thus reveals two properties-a highly asymmetrical, mainly negative PRC and second, a change of intrinsic period during entrainment.

### ERICA model and Arnold tongue

To gain understanding of these findings from a theory perspective, we next build a minimal model of our data.

First, we take the PRC computed at *T*_*zeit*_ = 140 mins to be the “reference” period, since it coincides with the natural period of the process. We can then estimate for each entrainment period the period shift most consistent with the data, to obtain a single, period independent PRC in Figure 8A2 (see Supplementary Note 2). From there, we build upon the simplest non-linear phase-oscillator, i.e. the classical Radial Isochron Cycle (RIC). To modulate its sine-form PRC, which is incompatible with our data, we perturb it into an Elliptic Radial Isochron Cycle with Acceleration or ERICA.

ERICA is designed to have radial isochrons, meaning that the phase (and the PRC) can be analytically computed in the entire plane. ERICA then allows for the speeding up of the cycle for some angle range (parameter *s*_*\**_) or for changes of the limit cycle shape into an ellipse of increasing eccentricity (parameter *λ*). Both high values of *λ* and *s*_*\**_ generate high negative asymmetry in the PRC (see Supplement; the full justification of the ERICA construction, the study of its multiple properties and its connection to biological oscillators will be described elsewhere). We then used Monte Carlo (MC) optimization to find parameters best fitting the experimental PRC. The results from this MC optimization put the oscillator far from the standard RIC oscillator (see cycle and corresponding flow in Figure 8B), consistent with strongly negative PRCs, with a moderate value of perturbation *A* = 0.4, high value of *λ* = 0.5 (indicating a strong elliptical shape), and high value of *s*_*\**_ = 5.6 over half a cycle (indicative of excitability). Figure 8C compares the PRC of this optimized model with the multiple data points, showing excellent agreement. In addition, we combined the ERICA framework with a simple fit for intrinsic period change to account for the experimental phase transition curves, as illustrated in Figure 8G.

With the ERICA model at hand, we derived numerically the Arnold tongues of the system and the phase/detuning curves for all entrainment parameters (Figure 8E-F). We notice that the Arnold tongues are heavily skewed toward the right, meaning that the system can be more easily slowed down than sped-up, consistent with the negative PRC shape. Remarkably, while we build the model using only one entrainment drug concentration, we can also largely explain data obtained at other concentrations (Figure 8F). In particular, the Arnold tongues/our model predict a specific change of the entrainment phase as the entrainment strength is varied (Figure S17G). The comparison to experimental data (Figure S10) shows this prediction is, qualitatively, verified. More generally, having the PRC and a minimal, coarse-grained model that captures the essential dynamical features of the segmentation clock during entrainment, including the change in intrinsic period, enables predictable control over the pace and the rhythm of the segmentation clock using the entrainment strategy.

## Discussion

In this work, we used a coarse-graining, entrainment approach to gain new insights into the dynamic properties of the segmentation clock from dynamical systems perspective. We mode-lock the segmentation clock to various entrainment periods and use the information about the dynamic phase-locking behaviour to derive the somite segmentation clock phase-response curve from the experimental data.

### A coarse-graining approach captures essential dynamical features using a simple “one oscillator” phase description

Given the complexity underlying the somite segmentation clock, comprised of several, interconnected, signaling pathways with countless molecular interactions, it was a priori unclear whether a simple “one oscillator” phase description and perturbation approach would capture the essential dynamical characteristics at the systems level. Another potential difficulty arises from the fact that cellular oscillators define a phase gradient, controlling the segmentation pattern (2). For all those reasons, one could imagine that no single phase could describe the systems behaviour, a general concern for systems of interacting oscillators already mentioned by Winfree (59).

Remarkably, one key finding of this work is that indeed, using a systems-level single oscillator phase description, we observe a consistent entrainment behaviour, i.e. period-locking with convergence towards a well-defined entrainment phase, as predicted from the oscillator phase response theory. It is important to point out that theoretical predictions are non-trivial and quantitative, such as the higher order 2 : 1 entrainment (Figure 5) and the dependence of the entrainment phase on detuning (Figure 7A-B). Given these experimental findings we conclude that the coarse-graining approach and the description of the entire system using a single oscillator phase is justified and enables to extract the essential dynamical properties, which are captured by a mathematical model including the systems’ PRC.

### Asymmetrical segmentation clock PRC compatible with saddle-node on invariant cycle (SNIC) bifurcation

Insight into the systems’ PRC allows to infer about the nature and characteristics of the oscillatory system, independent of its specific molecular realization (52). For instance, excitable systems can be functionally categorized with their PRC, i.e. in neuroscience, Class I excitable systems have PRCs with constant sign (52). The canonical example of (Class I) oscillators with constant PRCs is the quadratic fire and integrate model (52), which can be used to model neuronal spiking. In contrast, Class II excitable systems exhibit sinusoidal PRCs, examples include many biological systems such as the circadian clock (54).

This difference in PRC reflect fundamental properties of the underlying dynamics, including the associated bifurcations: Class I systems are associated to infinite period bifurcations, a typical example being the saddle-node on invariant cycle (SNIC) bifurcation. For such bifurcations, the period can be tuned and arbitrarily long depending on the proximity to the bifurcation, and oscillations present relaxation-type dynamics (i.e. combination of slow dynamics with fast resetting). The simplest example, the quadratic fire and integrate model, is a one variable *x* oscillator, where *x* goes from − ∞ to +∞ in a finite time before resetting to − ∞ (52). The PRC for this oscillator is “naturally” of constant sign because there is only one variable, which can only advance (or delay) in response to a constant signal. On the other hand, Class II systems are associated to Hopf bifurcations (52), have a fixed period and more sinusoidal dynamics close to the bifurcation.

Remarkably, the Segmentation Clock PRC that we computed here shares features of systems close to an infinite period bifurcation, i.e. a highly asymmetrical, constant sign PRC with an extended flat region. Consistent with this, the analytical ERICA model that we developed to capture the experimental PRCs in a wide range of conditions, exhibits relaxation-type dynamics (Fig. 8 B, slow phase in white and fast resetting phase in gray). It is nevertheless important to point out that a given PRC can result from different oscillators, i.e. a constant sign does not proof the existence of a SNIC oscillator. For instance, delayed oscillators close to a Hopf bifurcation can show PRCs with an (almost) constant sign (61). Infinite period can also be obtained with saddle homoclinic bifurcations (52). However such models are mathematically more complex, for instance delays include long-term memory in the system, and homoclinic bifurcations require the existence of other attractors outside of the cycle.

In agreement with SNIC oscillators, the segmentation clock does show a tunable period and a slowing down behaviour at multiple levels: it is known that the overall rate of segmentation slows down over developmental time, a feature described in several species. In addition, the oscillations also slow down along the embryonic axis, i.e. a period gradient is present within the PSM, reflecting tunability of oscillations at cellular level, as PSM cells progress towards differentiation and eventually stop oscillations. How the tunability at cellular level is linked to the system level change in segmentation clock rate is not understood (see also below for macro-vs. micro-scale comparison). However, our findings of mostly negative PRCs might reflect that the segmentation clock has a natural bias of slowing down the oscillations.

From a network standpoint, biological networks with tunable periods have been associated to specific structural features, combining negative and positive feedback loops (62). Positive feedback in particular puts oscillators close to a multistable regime, leading to relaxation oscillations (63). SNICs appear naturally when the regulatory logic of a system moves from a negative feedback oscillator to multistability (64), reflecting network modularity. Such modularity and robustness with changes in regulatory logic is a hallmark of developmental plasticity (65) and hence one would anticipate to find SNIC oscillators to be abundant in developing systems. First examples are indeed emerging, such as during *C. elegans* development (66).

### Findings not predicted by the theory of PRC

We also made several unexpected findings, not predicted by the theory of PRC

First, we found evidence that during entrainment, not only does the segmentation clock adjust its observed period to the *zeitgeber* pulses, but also, changes its intrinsic period in the direction of the *zeitgeber* rhythm. Hence, during entrainment that slows down the clock (i.e. 170 mins), we find evidence that the intrinsic period lengthens (not just the observed rhythm), while during entrainment that speeds up the clock (i.e. 120 mins), we find evidence that the intrinsic period decreased. Again, such a changes of intrinsic period is not predicted in entrained systems. The simplest explanation could be that the drug pulses change the period of the oscillator by changing some biochemical parameter in the system, similar to overdrive suppression in cardiac cells (60). However, as stated above, we do not find evidence for a consistent slowing down or speeding up effect: the intrinsic period is decreased for short entrainment cycles and increased for longer entrainment cycles. This rather suggests the existence of feedbacks of the clock on itself, leading to higher order adaptations beyond the rapid Notch phase response. One can only speculate on the mechanisms underlying such adaptation, but this is compatible with the idea that two interacting oscillators control the intrinsic period. Here, it is possible that entraining Notch with a *zeitgeber* modifies the Wnt oscillation period, which in turn feeds back on Notch on a longer time scale. So the internal period change that we see might in fact come from the induced change of Wnt period. We have shown previously Wnt and Notch oscillators are coupled but are not phase-locked, with functional impact on tissue patterning (1). Alternatively, a similar role could also be played by the long inter or intracellular delays in the system, postulated in multiple theoretical works (26, 67, 68). Such delays could effectively couple multiple cycles, changing clock parameters (such as the period) beyond instantaneous phase response. More experimental and theoretical work is needed to explore these ideas.

Second, a striking outcome we obtained was that even after entrainment, a spatial period gradient and phase waves emerged over time (Figure S11). Put differently, while the overall system is entrained, as evidenced by a control of the timing of morphological segmentation and of segmentation clock rhythm, the underlying cellular oscillators show a divergent, yet, coherent response, i.e. a spatial period gradient emerges within the tissue. A recent entrainment model of coupled PSM oscillators (69) does not account for the emergence of a period gradient during entrainment. This unexpected behaviour hence awaits further experimental and theoretical analysis. In particular, the role of intra- and intercellular coupling needs to be addressed to reveal how the macroscale behaviour relates to the underlying cellular scale oscillations.

## Conclusions

Our work demonstrates how, despite all the molecular and functional complexities, coarse-graining and theory can be used to effectively take control of complex biological processes. A *molecular* mechanism is not needed to exert control, as long as we have a *mathematical* one, one, that captures the essential features of a system at a meaningful coarse-grained level.

We also aim to illustrate the potential of an integrated, theoretical-experimental approach to complex biological systems: while from theoretical viewpoint, the fact that entrainment phase varies as detuning is altered is a mathematical necessity and hence ‘obvious’, this outcome is not at all intuitive, *a priori*, from an experimental-observational viewpoint. Theory guides experimentation, i.e. Figure 1 Theory, leading to new insight and understanding of a complex biological phenomenon.

Future studies are needed to gain further insight into the response to other perturbation regimes. In addition, it will be essential to perform the analysis both at the macro-scale, as done in this study, but also extending it to the cellular-scale, i.e. taking spatial differences, such as the period gradient, fully into account. We consider that such a dynamical systems theoretical-experimental approach is a promising and, as we show here, feasible way forward with the goal to categorize the segmentation clock in its universality class.

## Materials and methods

Please refer to supplementary materials (Supplementary Note 1) for detailed materials and methods.

## ACKNOWLEDGEMENTS

We thank Mogens Høgh Jensen, Jonas Juul, Mathias Heltberg, Sandeep Krishna, Adrián Granada and Istvan Kiss for in depth discussions on Arnold tongues and PRC; Justin Crocker, Anne Ephrussi, Lars Hufnagel, Jelle Scholtalbers, Ulrich Schwarz, Stefano Vianello, Nachiket Satish Shembekar and Ramesh Utharala, and all the members of the Aulehla group for the discussions and feedback. We are very grateful to Stefano Vianello for also designing some of the figures. We thank Antonio Politi for providing the Pipeline Constructor macro for automated live microscopy. We also acknowledge the assistance from Maximilian Beckers, Aliaksandr Halavatyi, and Oleksandr Maistrenko in writing scripts for data analysis. We are thankful to Ricardo Henriques for kindly sharing the template that was used to format this preprint. Following EMBL core facilities are acknowledged for their support of this work: the EMBL Advanced Light Microscopy Facility (ALMF), the EMBL Laboratory Animal Resources (LAR), the EMBL Mechanical Design Office and the EMBL Mechanical Workshop. The work was supported by the European Molecular Biology Laboratory (EMBL), by the German Research Foundation/DFG (CRC/SFB 1324, project number 331351713) to AA, the National Research Council of Canada, Discovery Grant program to Paul François, by the New Frontiers in Research Fund, Exploration program to PF and AA and the European Research Council (ERC) via an ERC Starting Grant (agreement n. 633943) and Consolidator Grant (agreement n.n.866537) to Alexander Aulehla.

## AUTHOR CONTRIBUTIONS

Paul Gerald Layague Sanchez designed the project, made the microfluidics devices, performed experiments, analyzed the data, and wrote the manuscript. Victoria Mochulska extracted PRCs, designed and optimized the ERICA model, and performed simulations. Christian Mauffette Denis optimized the model and performed Arnold tongue simulations. Gregor Mönke contributed to the project design and developed the wavelet analysis workflow used for the quantification of experimental time series. Takehito Tomita quantified and analyzed the period gradient. Nobuko Tsuchida-Straeten and Yvonne Petersen generated the Axin2-linker-Achilles mouse line. Katharina Sonnen developed the microfluidics-based experimental platform, performed experiments and contributed to the project design. Paul François designed the ERICA model, supervised the theory part of the project and wrote the manuscript. Alexander Aulehla designed and supervised the project and wrote the manuscript.

---

**Fig. S1.**
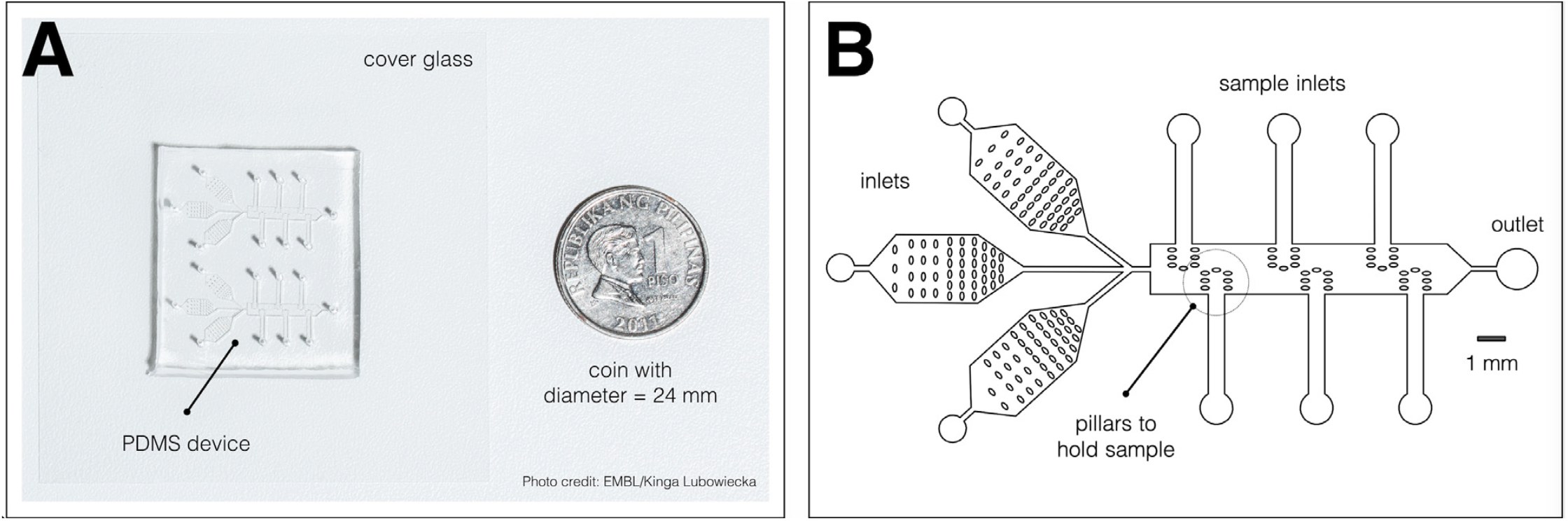
A microfluidics device for simultaneous culture, imaging, and entrainment of the segmentation clock in 2D-assays. (**A**) Photo of the chip, previously described in (1), bonded to cover glass and a coin (diameter: 24 mm) for scale. The split layout separating the upper and lower channel systems allows simultaneous delivery of drug and DMSO control to samples on opposite sides of the same device. Photo credit: EMBL/Kinga Lubowiecka. (**B**) Design of the microfluidics chip, showing inlets for medium and drug, inlets for the samples, pillars to hold each sample, and an outlet. A high-resolution version of this figure is available at https://doi.org/10.6084/m9.figshare.19093112.

**Table S1.** Summary statistics on period-locking of the segmentation clock in E10.5 2D-assays to periodic pulses of 2 uM DAPT. This table summarizes the median, 95% confidence interval (CI) of the median, mean, standard deviation (SD), and standard error of the mean (SEM) of the segmentation clock in 2D-assays subjected to periodic pulses of 2 uM DAPT. These summary statistics were determined using PlotsOfData (46). A plot of these data is shown in Figure 3D.

| Pulse Period | n | N | Median, mins | 95% CI of Median, mins | Mean, mins | SD, mins | SEM, mins |
| --- | --- | --- | --- | --- | --- | --- | --- |
| 120 mins | 14 | 3 | 127.36 | 123.16 – 148.03 | 139.39 | 22.28 | 6.18 |
| 130 mins | 39 | 10 | 133.44 | 132.21 – 135.97 | 134.75 | 4.56 | 0.74 |
| 140 mins | 15 | 3 | 147.68 | 146.10 – 153.02 | 150.00 | 7.42 | 1.98 |
| 150 mins | 17 | 4 | 155.14 | 154.33 – 160.74 | 158.18 | 7.77 | 1.94 |
| 160 mins | 15 | 3 | 167.09 | 166.16 – 169.84 | 167.33 | 5.28 | 1.41 |
| 170 mins | 34 | 8 | 176.14 | 173.06 – 181.20 | 175.56 | 8.52 | 1.48 |
| 180 mins | 6 | 1 | 184.43 | 164.54 – 194.21 | 181.06 | 17.88 | 8.00 |

**Fig. S2.**
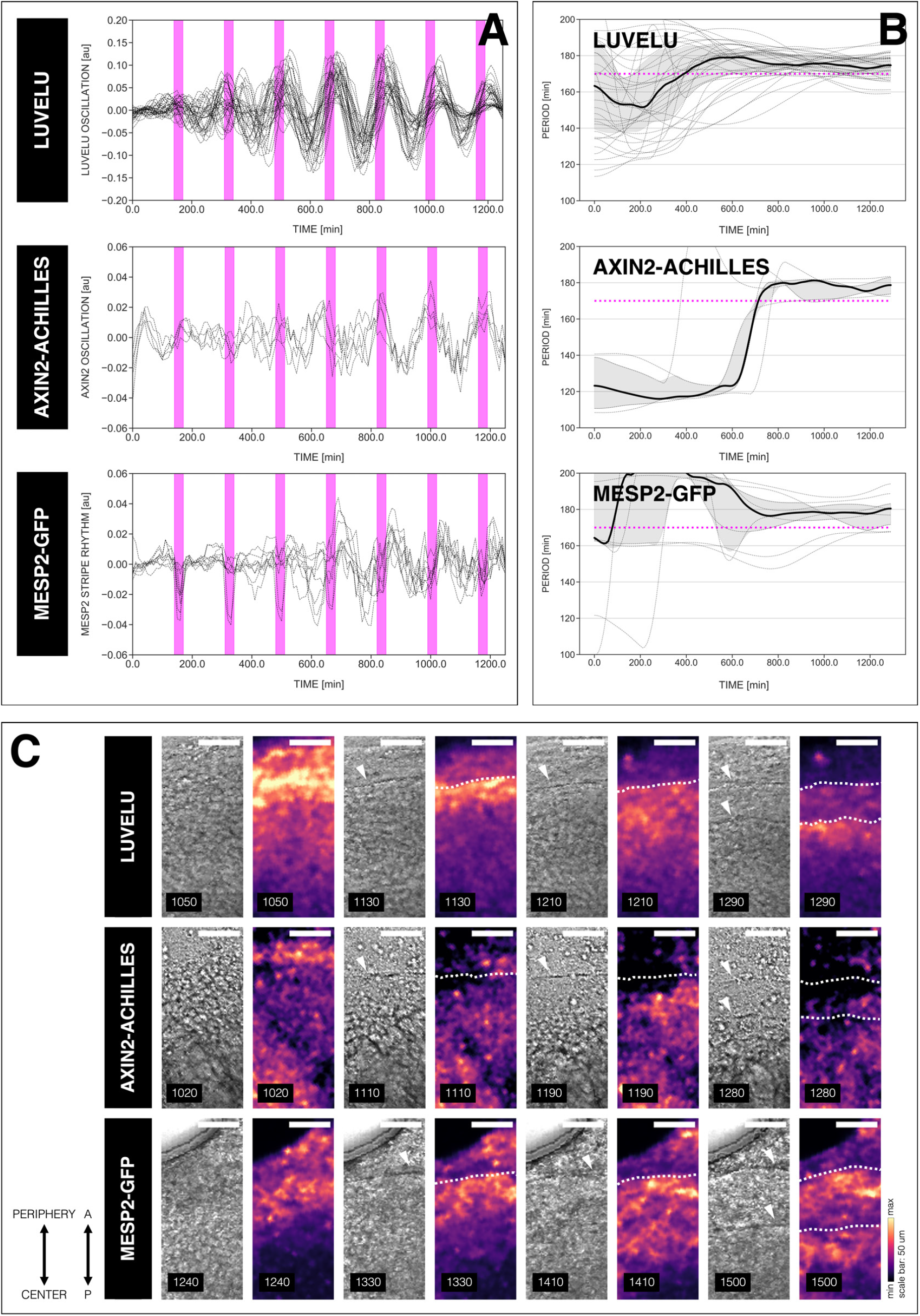
The rhythms of different dynamic readouts of the entrained segmentation clock match. (**A**) Detrended timeseries of the segmentation clock in 2D-assays subjected to 170-min periodic pulses of 2 uM DAPT, and expressing either LuVeLu (15), Axin2-linker-Achilles, or Mesp2-GFP (70). Periodic pulses are indicated as magenta bars and the timeseries of each sample (for LuVeLu: n = 34 and N = 8, for Axin2-linker-Achilles: n = 5 and N = 1, for Mesp2-GFP: n = 9 and N = 3) is marked with a dashed line. The detrended timeseries for samples expressing LuVeLu is the same as the detrended timeseries for the DAPT condition in Figure 3A. (**B**) Period evolution during entrainment, obtained from wavelet analysis. The period evolution for each sample and the median of the periods are represented here as a dashed line and a solid line, respectively. The gray shaded area corresponds to the interquartile range. Magenta dashed line marks *T*_*zeit*_. The period evolution plot for samples expressing LuVeLu is the same as the period evolution plot for the DAPT condition in Figure 3B. (**C**) Snapshots of segmenting regions in 2D-assays subjected to 170-min periodic pulses of 2 uM DAPT, and expressing either LuVeLu (15), Axin2-linker-Achilles, or Mesp2-GFP (70), at different time points. Segment boundaries are marked with either white arrowheads (brightfield channel) or white dashed lines (reporter channel). Samples are re-oriented so that the top is towards the periphery and the bottom is towards the center of the 2D-assay. Time is indicated as mins elapsed from the start of the experiment. Scale bar: 50 um. Timelapse movies of 2D-assays expressing either Axin2-linker-Achilles or Mesp2-GFP subjected to said perturbation are available at https://youtu.be/edFczx_-9hM and https://youtu.be/tQeBk0_U_Qo, respectively. A high-resolution version of this figure is available at https://doi.org/10.6084/m9.figshare.19093112.

**Fig. S3.**
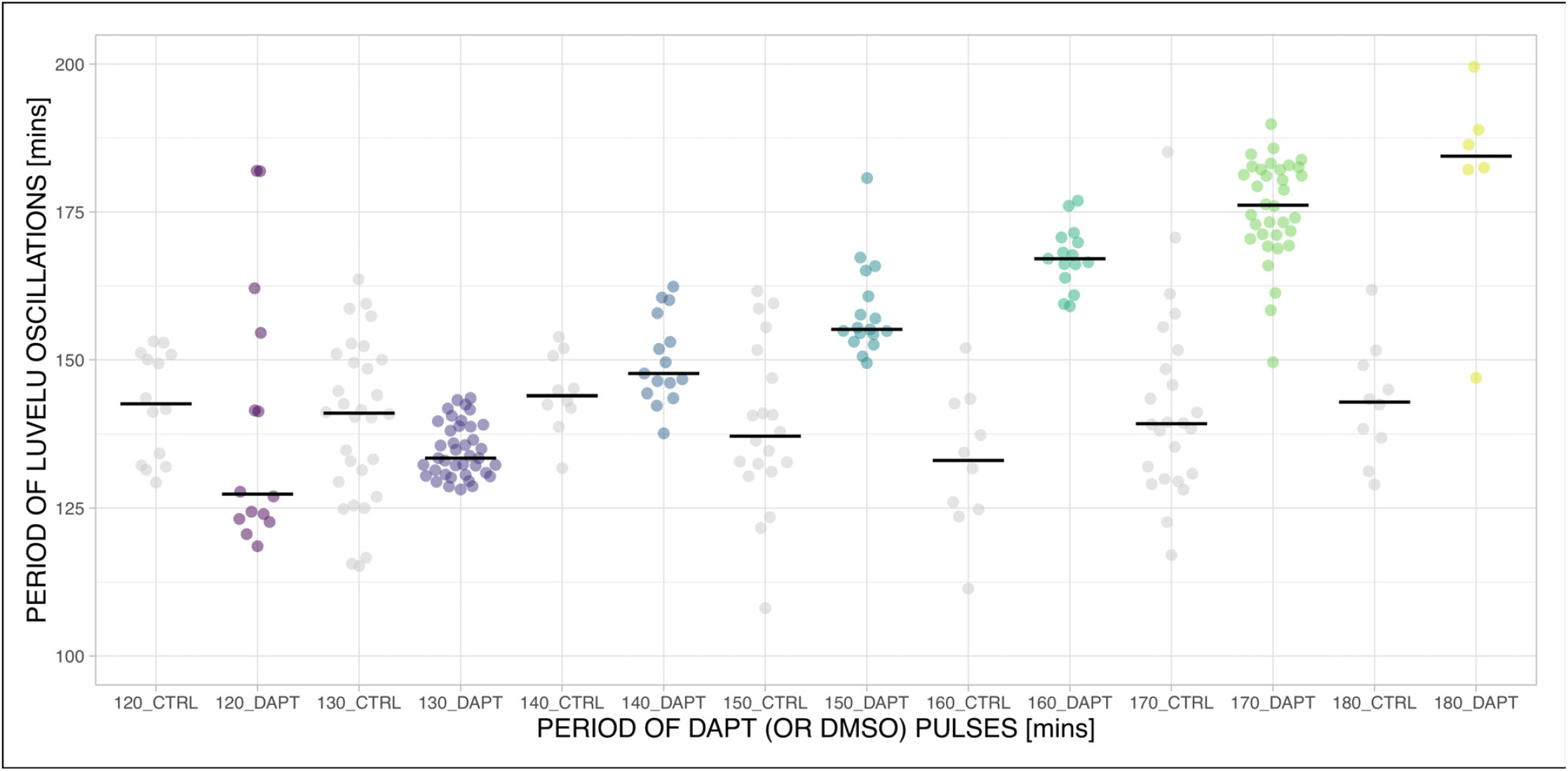
The period of the segmentation clock can both be sped up and slowed down by modulating the period of DAPT pulses. Mean period from 650 to 850 mins after start of the experiment of samples subjected to periodic pulses of 2 uM DAPT (or DMSO for controls). Each sample is represented as a dot, while the median of all samples is denoted as a solid horizontal line. The period of the DAPT (or DMSO) pulses is specified. **120-min**: (CTRL: n = 14 and N = 3) and (DAPT: n = 14 and N = 3) with p-value = 0.064, **130-min**: (CTRL: n = 30 and N = 8) and (DAPT: n = 39 and N = 10) with p-value = 0.003, **140-min**: (CTRL: n = 10 and N = 3) and (DAPT: n = 15 and N = 3) with p-value = 0.272, **150-min**: (CTRL: n = 20 and N = 4) and (DAPT: n = 17 and N = 4) with p-value = 0.001, **160-min**: (CTRL: n = 10 and N = 3) and (DAPT: n = 15 and N = 3) with p-value < 0.001, **170-min**: (CTRL: n = 24 and N = 7) and (DAPT: n = 34 and N = 8) with p-value < 0.001, **180-min**: (CTRL: n = 10 and N = 2) and (DAPT: n = 6 and N = 1) with p-value = 0.001. Data were visualized using PlotsOfData (46). To calculate p-values, two-tailed test for absolute difference between medians was done via a randomization method using PlotsOfDifferences (47). A high-resolution version of this figure is available at https://doi.org/10.6084/m9.figshare.19093112.

**Fig. S4.**
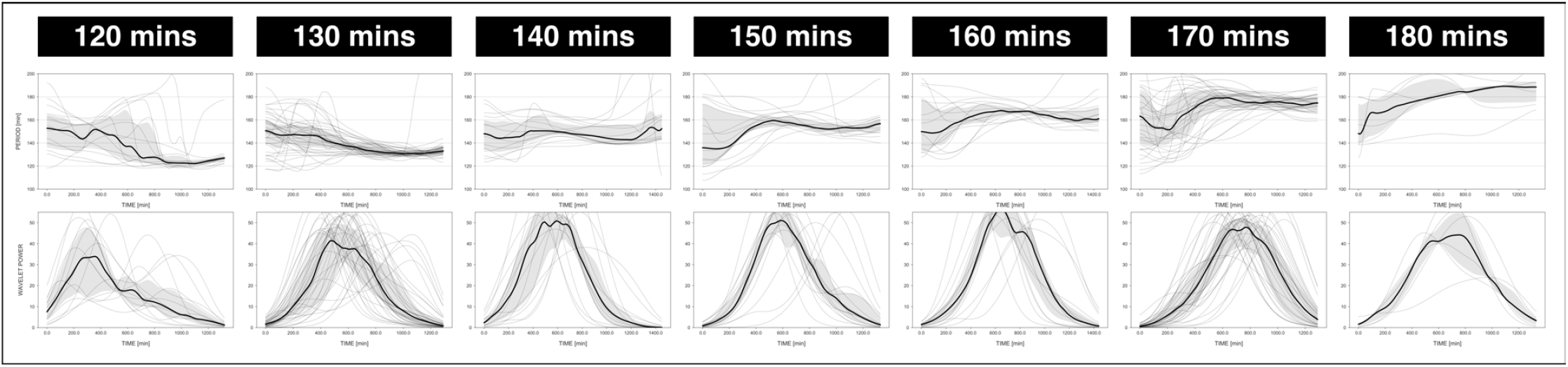
The period of the segmentation clock becomes locked to the period of the DAPT pulses. The period and wavelet power of the oscillations, obtained via wavelet analysis, are plotted across time. Each sample and their median are represented here as a dashed line and a solid line, respectively. The gray shaded area corresponds to the interquartile range. The period of the 2 uM DAPT pulses is specified. **120-min**: n = 14 and N = 3, **130-min**: n = 39 and N = 10, **140-min**: n = 15 and N = 3, **150-min**: n = 17 and N = 4, **160-min**: n = 15 and N = 3, **170-min**: n = 34 and N = 8, **180-min**: n = 6 and N = 1. The period evolution plots for the 130-min and 170-min conditions are the same as the period evolution plots for the 2 uM condition in Figure S5A and Figure 4B, respectively. A high-resolution version of this figure is available at https://doi.org/10.6084/m9.figshare.19093112.

**Fig. S5.**
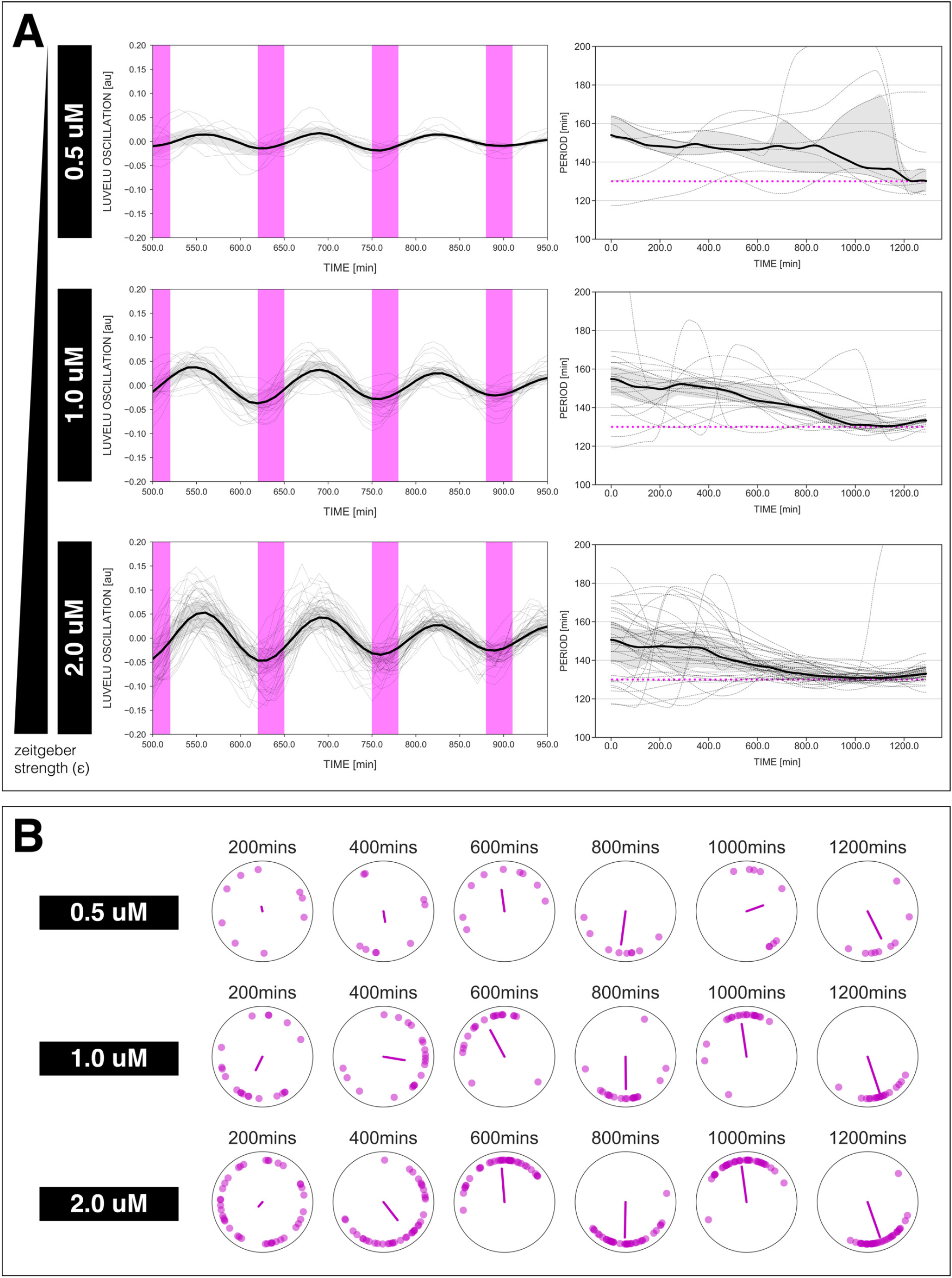
Changing the concentration of DAPT, equivalent to changing *zeitgeber* strength, affects entrainment of the segmentation clock to 130-min periodic DAPT pulses. (**A**) Left: Detrended (via sinc-filter detrending) timeseries of the segmentation clock in 2D-assays entrained to 130-min periodic pulses of either 0.5 uM, 1 uM, or 2 uM DAPT, zoomed in from 500 mins to 950 mins. Periodic pulses are indicated as magenta bars and the timeseries of each sample (for 0.5 uM: n = 9 and N = 2, for 1 uM: n = 20 and N = 4, for 2 uM: n = 39 and N = 10) is marked with a dashed line. The median of the oscillations is represented here as a solid line, while the gray shaded area denotes the interquartile range. Right: Period evolution during entrainment, obtained from wavelet analysis. The period evolution for each sample and the median of the periods are represented here as a dashed line and a solid line, respectively. The gray shaded area corresponds to the interquartile range. Magenta dashed line marks *T*_*zeit*_. (**B**) Polar plots at different timepoints showing phase of each sample and their first Kuramoto order parameter, represented as a magenta dot along the circumference of a circle and a magenta line segment at the circle’s center, respectively. A longer line segment corresponds to a higher first Kuramoto order parameter, and thus to more coherent samples. The direction of the line denotes the vectorial average of the sample phases. Time is indicated as mins elapsed from the start of the experiment. A high-resolution version of this figure is available at https://doi.org/10.6084/m9.figshare.19093112.

**Fig. S6.**
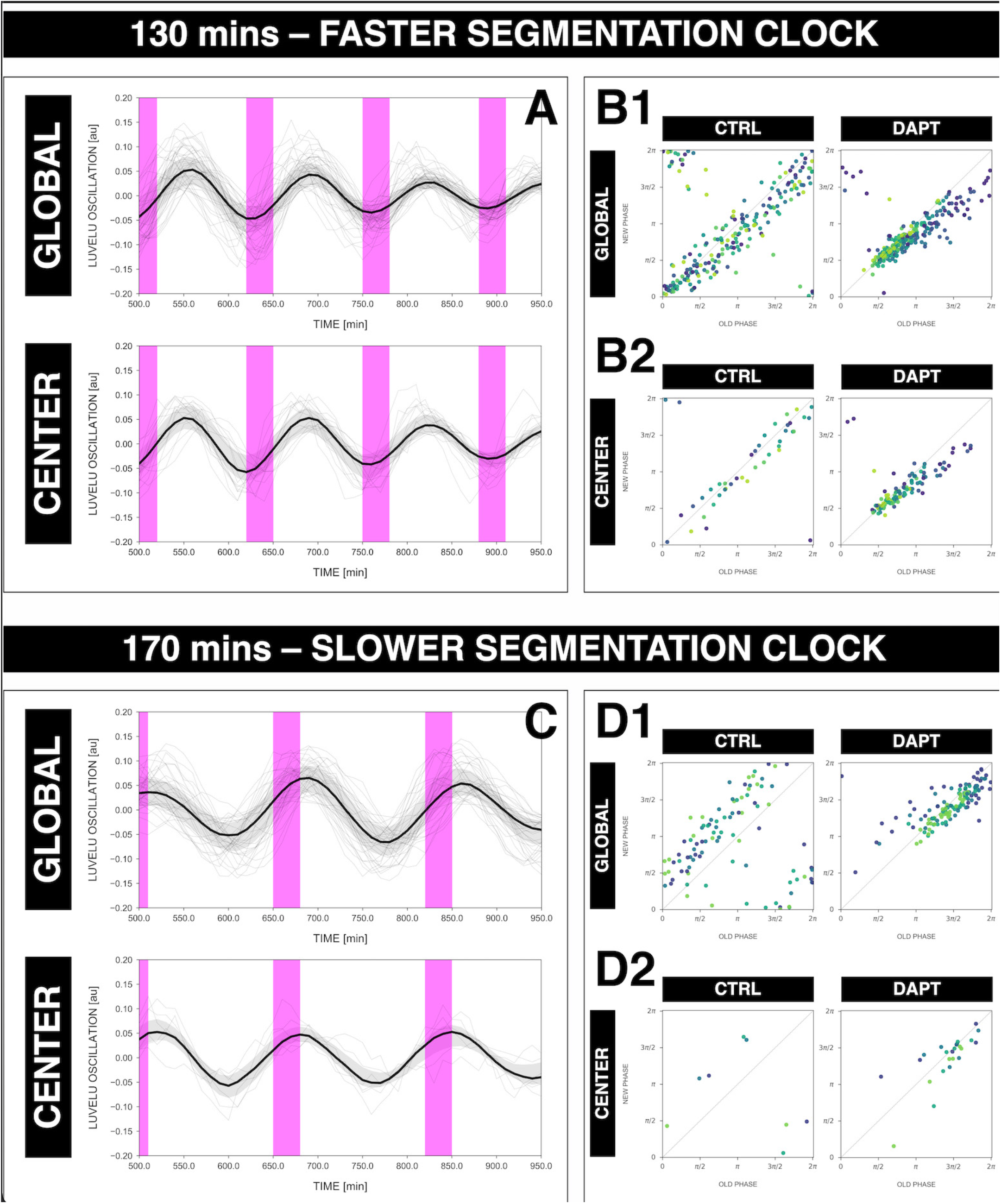
The rhythm of the segmentation clock matches the rhythm of oscillations in the center of 2D-assays (i.e. the posterior PSM) regardless whether the clock is sped up or slowed down. (**A**) Detrended timeseries of either the segmentation clock (global ROI, ‘GLOBAL’) or oscillations in the center of 2D-assays (‘CENTER’) entrained to 130-min periodic pulses of 2 uM DAPT, zoomed in from 500 mins to 950 mins. Periodic pulses are indicated as magenta bars and the timeseries of each sample (for GLOBAL: n = 39 and N = 10, for CENTER: n = 15 and N = 7) is marked with a dashed line. The median of the oscillations is represented here as a solid line, while the gray shaded area denotes the interquartile range. The plot for GLOBAL is the same as that for the 130-min condition in Figure 7C. (**B**) Stroboscopic maps of either the segmentation clock (GLOBAL, B1) or oscillations in the center of 2D-assays (CENTER, B2) subjected to 130-min periodic pulses of 2 uM DAPT, with their respective controls (subjected to periodic pulses of DMSO). Colors mark progression in time, from purple to yellow. (**C**) Detrended timeseries (via sinc-filter detrending) of either the segmentation clock (GLOBAL) or oscillations in the center of 2D-assays (CENTER) entrained to 170-min periodic pulses of 2 uM DAPT, zoomed in from 500 mins to 950 mins. Periodic pulses are indicated as magenta bars and the timeseries of each sample (for GLOBAL: n = 34 and N = 8, for CENTER: n = 6 and N = 5) is marked with a dashed line. The median of the oscillations is represented here as a solid line, while the gray shaded area denotes the interquartile range. The plot for GLOBAL is the same as that for the 170-min condition in Figure 7C. (**D**) Stroboscopic maps of either the segmentation clock (GLOBAL, D1) or oscillations in the center of 2D-assays (CENTER, D2) subjected to 170-min periodic pulses of 2 uM DAPT, with their respective controls (subjected to periodic pulses of DMSO). Colors mark progression in time, from purple to yellow. A high-resolution version of this figure is available at https://doi.org/10.6084/m9.figshare.19093112.

**Fig. S7.**
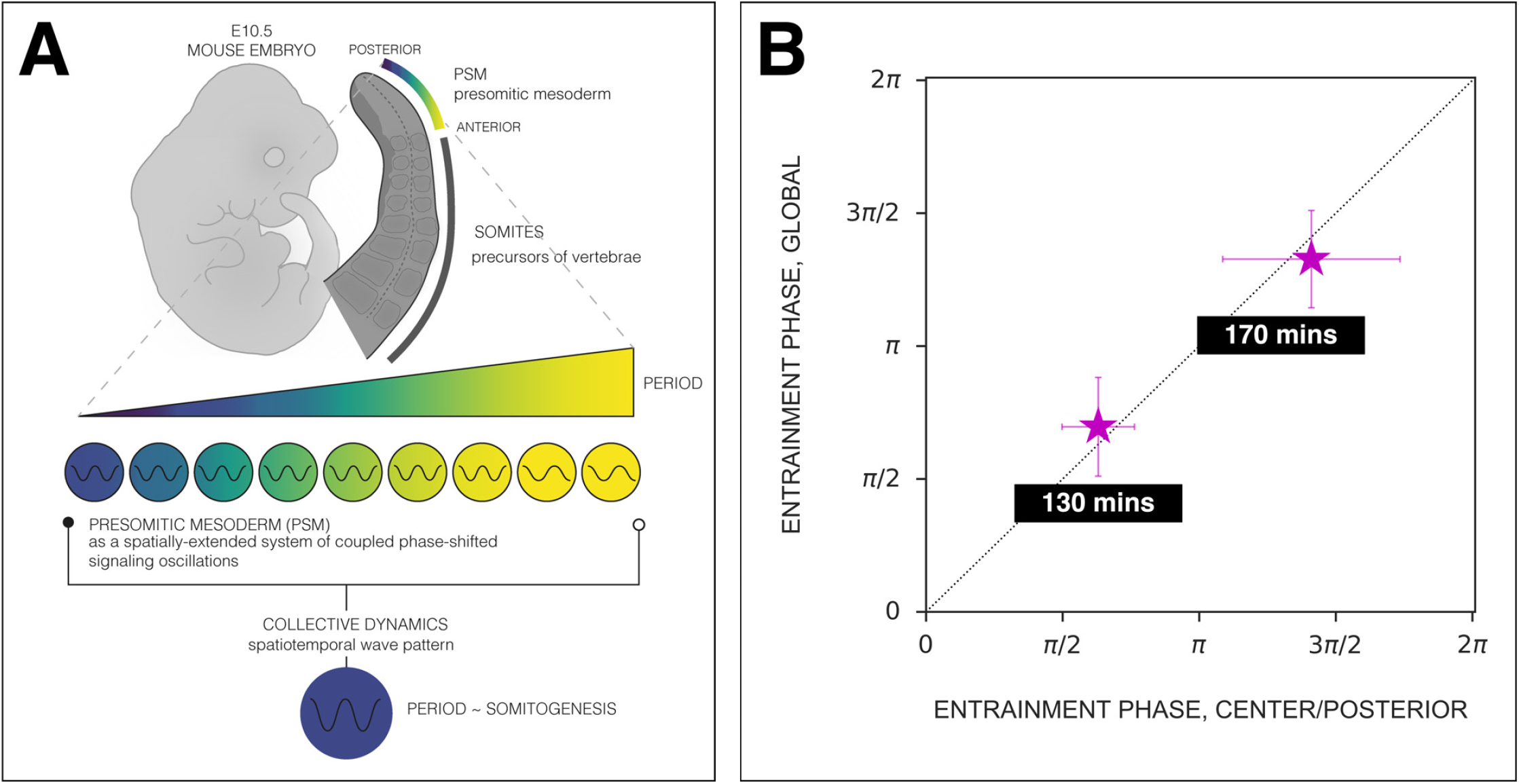
The phase of oscillations in the the center of 2D-assays(posterior PSM) matches the phase of the segmentation clock. (**A**) Scheme of the phase-shifted oscillations along the AP axis of the PSM and its correlation to the segmentation clock. Illustration by Stefano Vianello. (**B**) Comparison of the phase relationship between the periodic DAPT pulses and either the segmentation clock rhythm (measured in global ROI, ‘GLOBAL’) or the rhythm in the center of 2D-assays (equivalent to posterior PSM, CENTER/POSTERIOR). The entrainment phase was calculated from the vectorial average of the phases of all samples at the time corresponding to the last DAPT pulse considered. The spread of sample phases is reported in terms of the circular standard deviation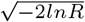, where R is the first Kuramoto order parameter). **130-min**: (GLOBAL: n = 39 and N = 10) and (CENTER: n = 15 and N = 7), **170-min**: (GLOBAL: n = 34 and N = 8) and (CENTER: n = 6 and N = 5). A high-resolution version of this figure is available at https://doi.org/10.6084/m9.figshare.19093112.

**Fig. S8.**
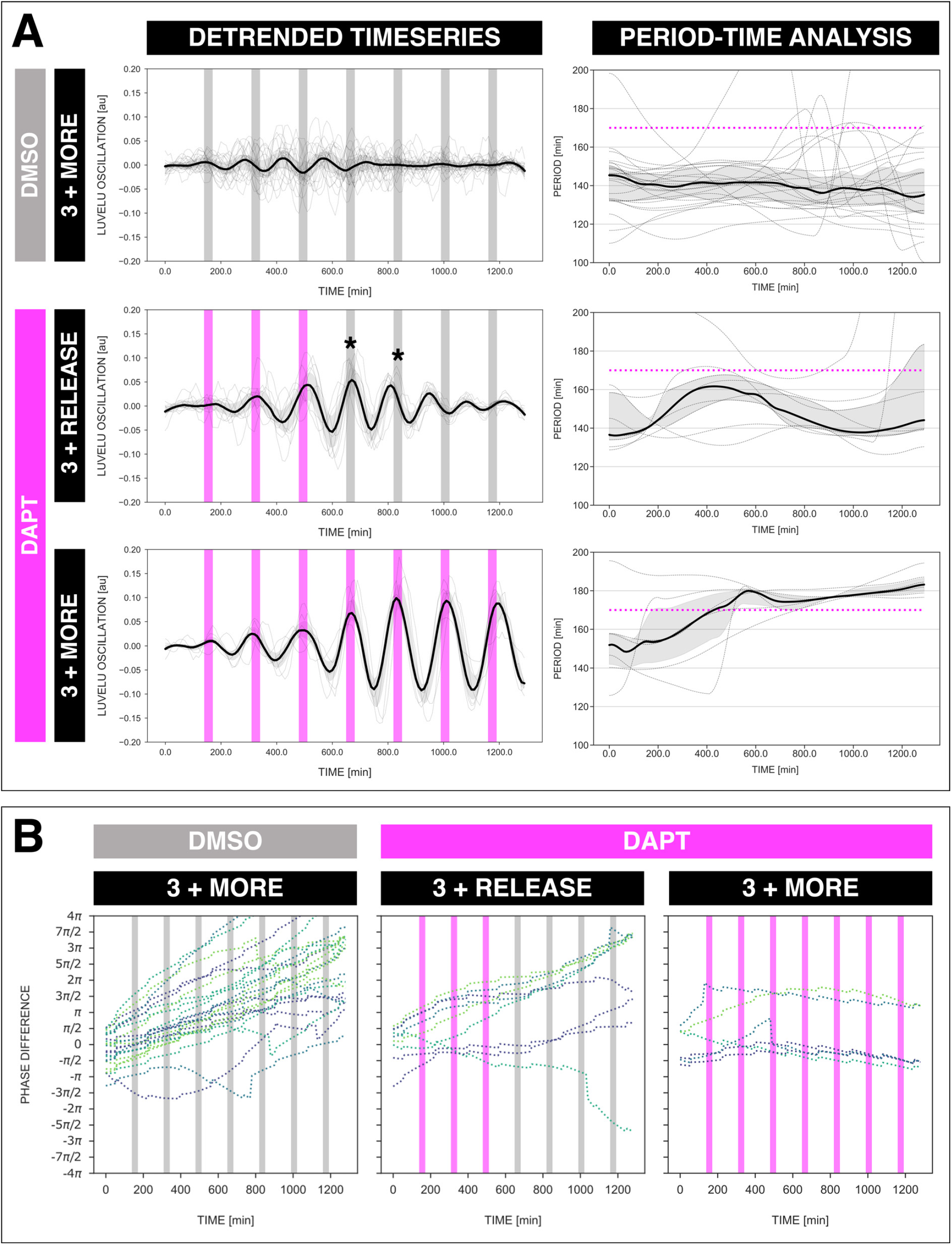
The segmentation clock keeps its adjusted rhythm even a few cycles after release from DAPT pulses. (**A**) Left: Detrended (via sinc-filter detrending) timeseries of the segmentation clock in 2D-assays subjected to 170-min periodic pulses of DMSO (gray bars) and/or 2 uM DAPT (magenta bars). The timeseries of each sample (for continuous DMSO pulses: n = 24 and N = 7, for 3 DAPT pulses and then release: n = 9 and N = 2, for continuous DAPT pulses: n = 6 and N = 2) is marked with a dashed line. The median of the oscillations is represented here as a solid line, while the gray shaded area denotes the interquartile range. Right: Period evolution during entrainment, obtained from wavelet analysis. The period evolution for each sample and the median of the periods are represented here as a dashed line and a solid line, respectively. The gray shaded area corresponds to the interquartile range. Magenta dashed line marks *T*_*zeit*_. Data for the continuous DMSO pulses are the same as the controls in Figure 3A-B. (**B**) Phase difference between the segmentation clock and the drug pulses. Note that a phase of -*π*/2 is equivalent to a phase of 3*π*/2, and a phase of 0 is equivalent to a phase of 2*π*. Periodic pulses of DMSO and 2 uM DAPT are indicated as gray bars and as magenta bars, respectively. Each sample within each condition is marked with different colors. A high-resolution version of this figure is available at https://doi.org/10.6084/m9.figshare.19093112.

**Fig. S9.**
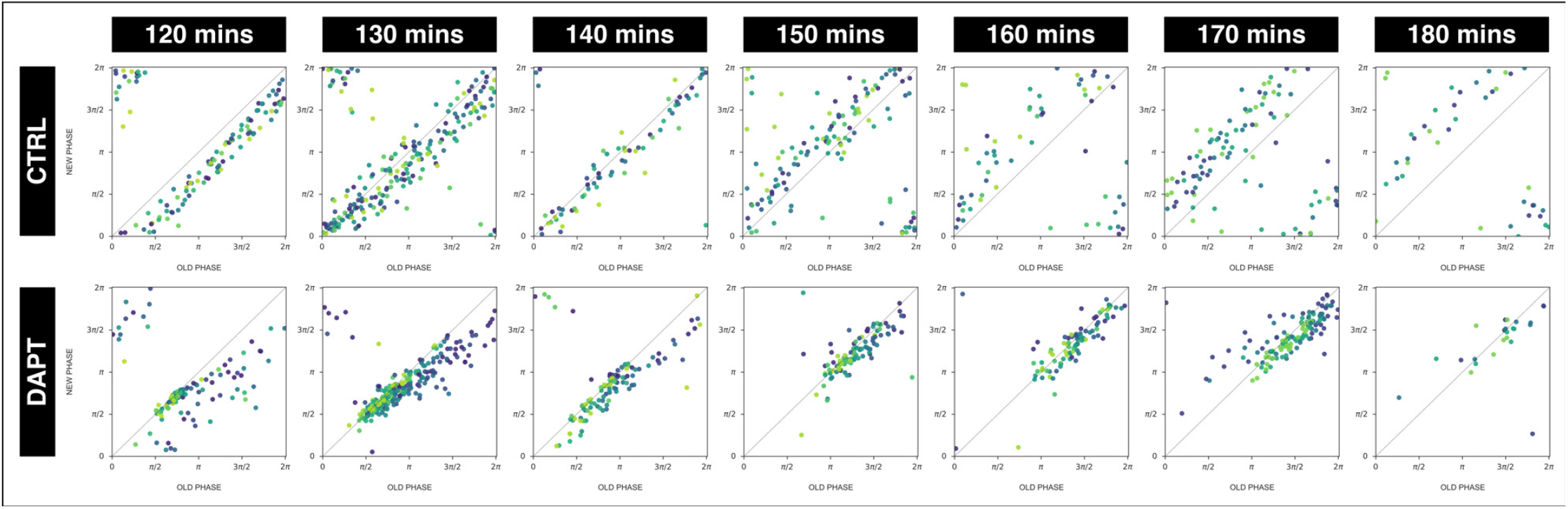
The segmentation clock establishes a stable phase relationship with the periodic DAPT pulses. Stroboscopic maps summarizing phase dynamics when the segmentation clock was subjected to periodic pulses of 2 uM DAPT (or DMSO for controls). The period of the DAPT (or DMSO) pulses is specified. Colors mark progression in time, from purple to yellow. **120-min**: (CTRL: n = 14 and N = 3) and (DAPT: n = 14 and N = 3), **130-min**: (CTRL: n = 30 and N = 8) and (DAPT: n = 39 and N = 10), **140-min**: (CTRL: n = 10 and N = 3) and (DAPT: n = 15 and N = 3), **150-min**: (CTRL: n = 20 and N = 4) and (DAPT: n = 17 and N = 4), **160-min**: (CTRL: n = 10 and N = 3) and (DAPT: n = 15 and N = 3), **170-min**: (CTRL: n = 24 and N = 7) and (DAPT: n = 34 and N = 8), **180-min**: (CTRL: n = 10 and N = 2) and (DAPT: n = 6 and N = 1). A high-resolution version of this figure is available at https://doi.org/10.6084/m9.figshare.19093112.

**Fig. S10.**
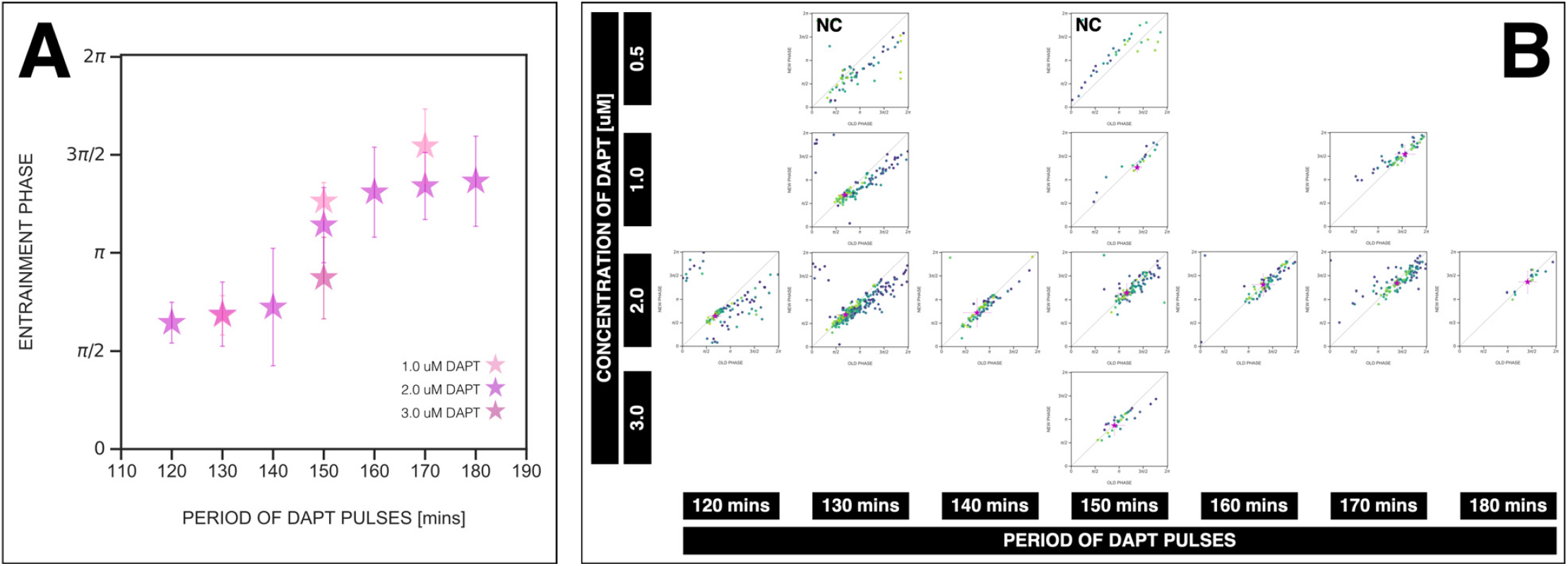
*Zeitgeber* period and *zeitgeber* strength affect the entrainment phase of the segmentation clock. (**A**) Entrainment phase at different periods of DAPT pulses (i.e. *zeitgeber* period) and different drug concentrations (i.e. *zeitgeber* strength). Entrainment phase (*φ*_*ent*_) was calculated from the vectorial average of the phases of phase-locked samples at the time corresponding to last considered DAPT pulse. A sample was considered phase-locked if the difference between its phase at the time of the final drug pulse considered and its phase one drug pulse before is less than *π/*8 – for 120-min: (2 uM: n = 13/14 and N = 3/3), for 130-min: (1 uM: n = 17/20 and N = 4/4), (2 uM: n = 38/39 and N = 10/10), for 140-min: (2 uM: n = 10/15 and N = 3/3), for 150-min: (1 uM: n = 3/4 and N = 1/1), (2 uM: n = 16/17 and N = 4/4), (3 uM: n = 5/5 and N = 1/1), for 160-min: (2 uM: n = 11/15 and N = 3/3), for 170-min: (1 uM: n = 12/18 and N = 4/5), (2 uM: n = 28/34 and N = 8/8), for 180-min: (2 uM: n = 4/6 and N = 1/1). The spread of *φ*_*ent*_ between samples is reported in terms of the circular standard deviation 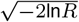, where *R* is the first Kuramoto order parameter). Colors mark concentration of DAPT. Drug pulse duration was kept constant at 30 mins/cycle. (**B**) Stroboscopic maps for different values of *zeitgeber* period and *zeitgeber* strength placed next to each other. The localized region close to the diagonal in each map marks *φ*_*ent*_ for that condition. This is highlighted with a magenta star, which corresponds to the centroid of the said region. The centroid (*x*_*c*_, *y*_*c*_) was calculated from the vectorial average of the phases of phase-locked samples at the end of the experiment, where *x*_*c*_ = vectorial average of old phase, *y*_*c*_ = vectorial average of new phase. The spread of the points in the region is reported in terms of the circular standard deviation 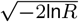, where *R* is the first Kuramoto order parameter). The period of the DAPT pulses and the concentration of DAPT are indicated. Colors mark progression in time, from purple to yellow. NC means not considered (in plotting panel A) because no/small fraction of samples was phase-locked – for 0.5 uM 130-min: n = 4/9 and N = 2/2, for 0.5 uM 150-min: n = 0/6 and N = 0/1. Stroboscopic maps for these NC conditions include all samples, unlike the other conditions where only phase-locked samples are plotted. Data for 2 uM condition is the same as that in Figure 7A-B. A high-resolution version of this figure is available at https://doi.org/10.6084/m9.figshare.19093112.

**Fig. S11.**
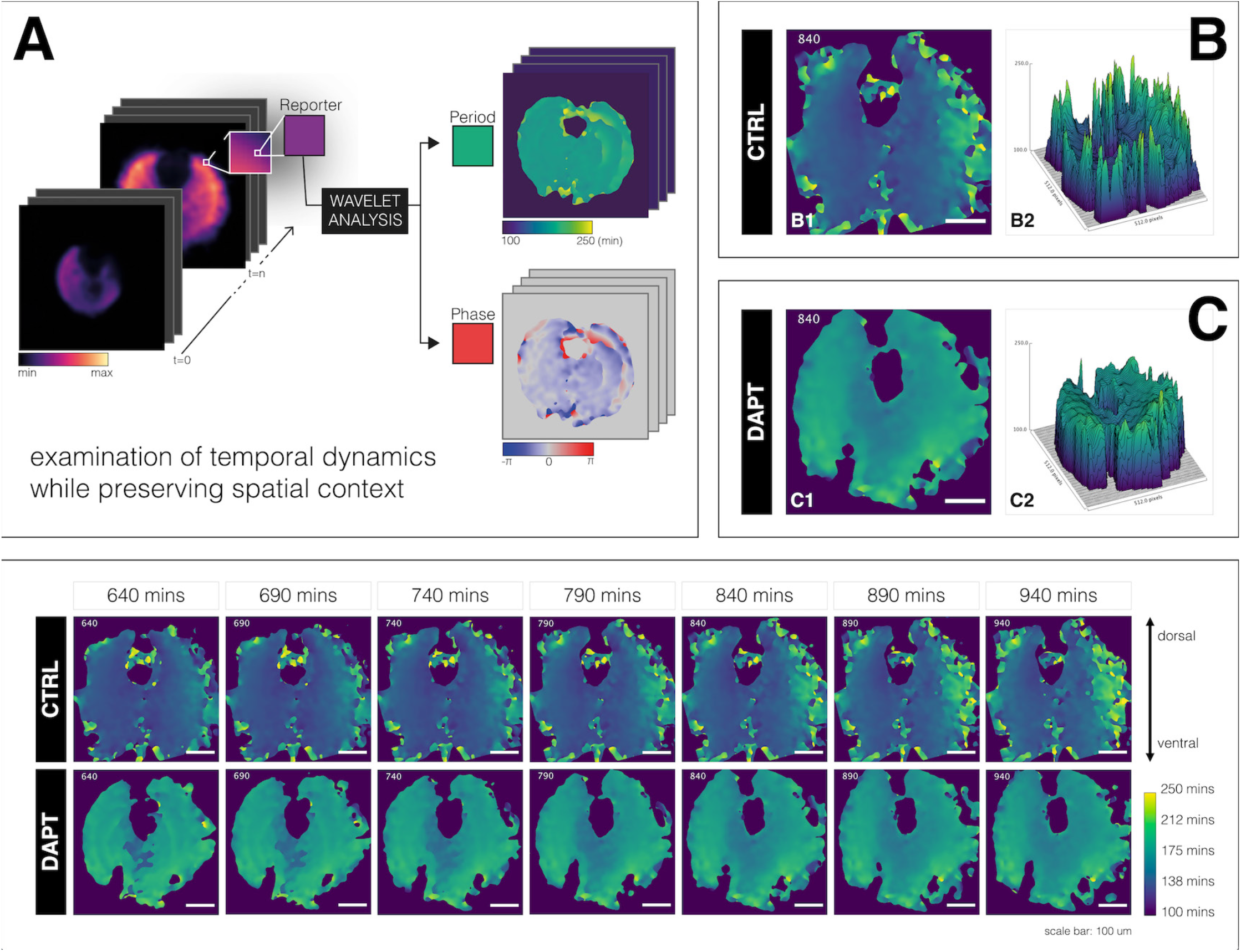
A spatial period gradient emerges in the tissue even upon entrainment of the segmentation clock. Left: Schematic of pipeline to generate period and phase movies using pixel-by-pixel wavelet analysis (**A**). Illustration by Stefano Vianello. **Right**: Snapshot of period wavelet movie of control subjected to 170-min periodic pulses of DMSO (**B**) or entrained sample subjected to 170-min periodic pulses of 2 uM DAPT (**C**), taken at 840 mins after the start of experiment. Period is either shown using heatmap (B1,C1) or as a surface plot (B2,C2). **Bottom**: Snapshots of period wavelet movie of samples in (B) and (C) at different time points. Time is indicated as mins elapsed from the start of the experiment. Sample is rotated so that the dorsal side is up. A high-resolution version of this figure is available at https://doi.org/10.6084/m9.figshare.19093112. Timelapse of period wavelet movies and corresponding surface plots are available at https://youtu.be/3Y53OhXacKI.

## Supplementary Note 1: Materials and methods

### A. Mouse lines. [0.2em]

For most of the experiments in this study, we used a transgenic mouse line expressing a dynamic Notch signaling reporter driven from the Lfng promoter, more commonly known as LuVeLu. The generation of LuVeLu was previously described (15). Briefly, the expression of Venus, an improved version of YFP (71), was driven from a 2-kb fragment of the Lfng promoter (72, 73). The locus was flanked by Lfng 3’-UTR and a modified PEST domain (74) to destabilize the reporter mRNA and protein, respectively. For experiments visualizing Mesp2 during entrainment, we used the Mesp2-GFP mouse line generated by (70).

Axin2-linker-Achilles, a mouse line expressing Axin2 fused with a GSAGS linker to Achilles, a fast-maturing variant of YFP (23), was generated in-house. To generate the knock-in alleles, we targeted the stop codon of endogenous Axin2 locus with vector containing the reporter sequence coding for Achilles and a selection cassette. This targeting vector was constructed as follows: linker-Achilles-loxP-PGK Neo-loxP. The selection cassette was flanked by loxP-sites for eventual Cre-mediated excision. Axin2-linker-Achilles knock-in reporter line was generated by standard gene targeting techniques using R1 embryonic stem cells. Briefly, chimeric mice were obtained by C57BL/6 blastocyst injection and then outbred to establish the line through germline transmission. The Achilles/pRSETB plasmid was a gift from the lab of Atsushi Miyawaki at RIKEN Center for Brain Science (RIKEN-CBS) in Japan.

Mice were kept in an outbred background and were housed in the EMBL Laboratory Animal Resources (LAR). All animal experiments were conducted under veterinarian supervision and after project approval by European Molecular Biology Laboratory, following the guidelines of the European Commission, Directive 2010/63/EU and AVMA Guidelines 2007.

### B. Media preparation

On the day of the experiment, dissection medium and culture medium were freshly prepared as indicated in Table S2. Culture medium was filter sterilized using a PVDF filter (pore size: 0.22 um, Merck). Both dissection medium and culture medium were equilibriated to 37^*°*^C for at least 15 minutes, and were kept in a 37^*°*^C incubator under 5% CO_2_ until use.

**Table S2.**
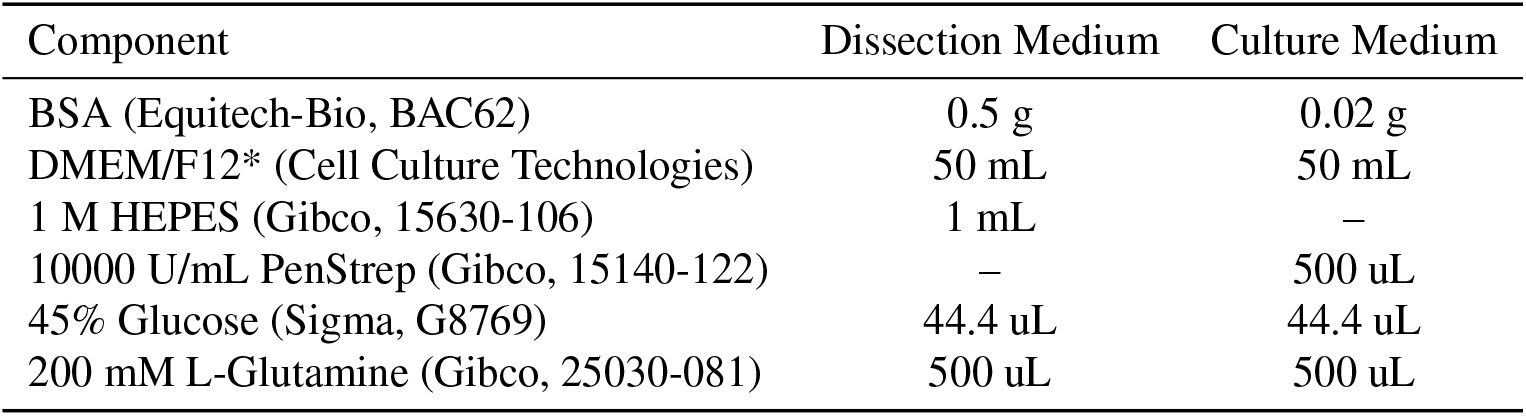
Recipe for media preparation. Formulations specified here are for the preparation of approximately 50 mL of medium. Special DMEM/F12* used in these media does not contain glucose, L-glutamine, sodium pyruvate, and phenol red. Culture medium is filter sterilized after preparation.

### C. Mouse dissection and embryo recovery

Female mice were sacrificed on 10.5 dpc (days post coitum) via cervical dislocation. The skin on their ventral side (belly area) was wiped with 70% ethanol, and an incision was made using a clean pair of surgical scissors. The uterine horns were harvested and were washed once with dissection medium. In dissection medium, under a stereo microscope (Leica M80), the deciduae were cut open using clean forceps and the embryos were recovered. The embryos were again washed with fresh dissection medium and their tails were clipped using forceps. Clipped tails were then transferred to a new dish with fresh dissection medium. The volume of dissection medium in the dish was kept to minimum to lessen autofluorescence, which could interfere with subsequent screening. The tails were then screened for presence of the reporter-of-interest (e.g. LuVeLu) using a stereo fluorescence microscope.

The tailbud was cut from the rest of the tail and was immediately transferred to pre-equilibriated culture medium that was supplemented with HEPES (170 uL of 1 M HEPES in 10 mL culture medium). These isolated embryonic tissues were then directly loaded in the microfluidics device.

### D. Preparations for microfluidics-based entrainment of 2D-assays

We used in this study a microfluidics-based experimental entrainment platform with a general protocol elaborated in a recent publication (75). The PDMS microfluidics device was made using standard soft lithography (76) and as previously described (1). The ratio of Sylgard 184 silicone elastomer base (Dow) to curing agent (Dow) was 9:1 (w/w). PDMS chip was attached to cover glass (70 mm x 70 mm, 1.5H, Marienfeld 0107999 098) via plasma bonding.

#### UV irradiation of PTFE tubing and PDMS device

PTFE tubing (inner diameter: 0.6 mm, APT AWG24T) and syringe needles (22G 1 1/4 - Nr. 12), as summarized in Table S3, were prepared a day prior to actual microfluidics-based entrainment experiment. In addition to 1 PDMS microfluidics device (Figure S1B), each experiment required 4 3-meter PTFE tubing (each with syringe needle inserted in one end) for the drug/medium inlets, 2 1-meter PTFE tubing (each with syringe needle inserted in one end) for the outlets, and 24 1-centimeter plugs made from cut PDMS-filled PTFE tubing for the sample inlets and unused drug/medium inlets. These were all sterilized under UV for at least 20 minutes.

**Table S3.** List of PTFE tubing needed for a microfluidics-based entrainment experiment. For the first two items, a syringe needle is to be inserted inside one end of each tubing. Plugs are made from cut PDMS-filled PTFE tubing, and are used to seal inlets after sample loading. Controls are already taken into account in the specified quantities.

| Item | With Needle? | Quantity | Use |
| --- | --- | --- | --- |
| 3-meter PTFE tubing | Y | 4 | drug/medium inlet |
| 1-meter PTFE tubing | Y | 2 | outlet |
| 1-centimeter plug | N | 24 | sample inlet + unused drug/medium inlet |

#### Fibronectin-coating of PDMS device and overnight soaking in buffer

While waiting, 5 mL of PenStrep (Gibco, 15140-122) was added to 495 mL of 1x PBS (PBS+PenStrep). 5.6 mL of the buffer was set aside to prepare fibronectin solution, while the rest was poured in a glass dish. After UV irradiation, the sterilized PDMS device was immersed in the PBS+PenStrep and bubbles were removed by flushing the channels with buffer. PDMS-filled PTFE plugs were also immersed in PBS+PenStrep. To prepare the fibronectin solution, 280 uL of fibronectin (Sigma-Aldrich, F1141) was added to the set aside 5.6 mL PBS+PenStrep. At least 2.5 mL of fibronectin solution was loaded into a 3 mL syringe (diameter: 8.66 mm, BD Luer-Lok REF 309658). A UV-irradiated needle, which was earlier inserted into a 1-meter PTFE tubing, was attached to the filled syringe. The tubing was then inserted into an outlet in the PDMS device, carefully avoiding introduction of bubbles. Fibronectin was flowed (flow rate: 50 uL/hr) into the PDMS device at room temperature overnight.

#### Preparation of syringes containing drug/medium

For a microfluidics-based experiment with periodic pulses of drug, four 10 mL syringes (diameter: 14.5 mm, BD Luer-Lok REF 300912) were filled with either the drug, DMSO control, or culture medium (see Table S2). Components of solution in each of these syringes are specified in Table S4. Drug used in this study was DAPT (Sigma-Aldrich, D5942-5MG). To prepare 10 mM stock of DAPT, 5 mg DAPT (MW = 432.46 g/mol) was dissolved in 1156.2 uL DMSO (Sigma-Aldrich, D8418). This was aliquoted and stored at -20^*°*^C until use.

**Table S4.**
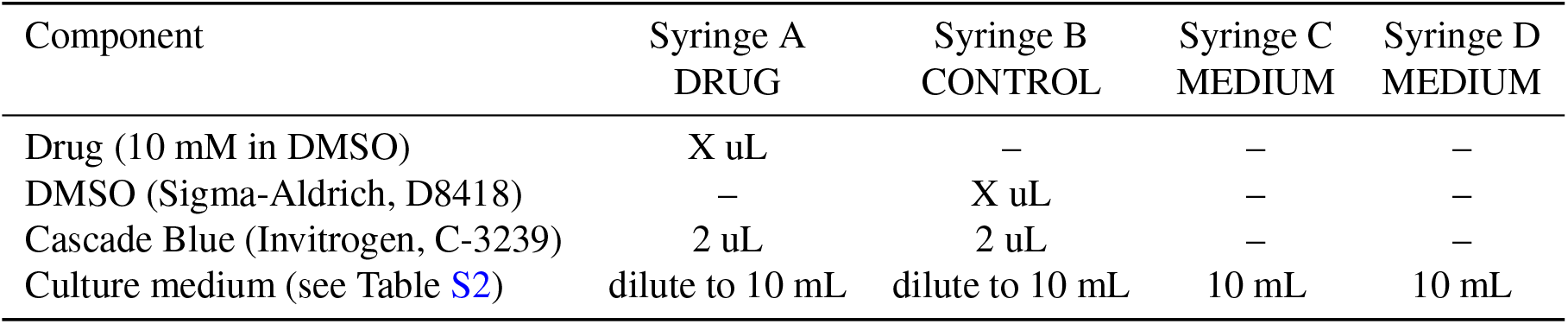
List of syringe containing drug/medium for a microfluidics-based entrainment experiment. Formulations specified here are for any drug with final concentration X uM. For recipe to prepare culture medium, please refer to Table S2.

#### Degassing drug/medium and PDMS device

After coating and overnight soaking, the PTFE tubing was cut away from the needle and was immediately immersed in the buffer. The dish containing immersed PDMS device, with attached tubing for the two outlets, and plugs were placed inside a vacuum desiccator chamber. The plunger of each syringe containing the drug/medium was pulled to maximum. The syringes were then also placed in the vacuum chamber, almost vertically, with the plunger resting on the desiccator. These were degassed under high pressure for at least 1.5 hours.

### E. Loading embryonic tissues in microfluidics device and mounting for live imaging

Before mouse dissection and recovery of mouse embryos, syringes containing the degassed drug/medium were each connected to a UV-irradiated 3-meter PTFE tubing via attached syringe needle, and were carefully mounted on programmable pumps (World Precision Instruments, AL-400) next to the microscope. Gas in the tubing was displaced with drug/medium by careful pushing of the syringes’ plunger. Pumps were turned on and flow rate was set to 900 uL/hr. The microscope was then equilibriated to 37^*°*^C and 5% CO_2_.

After recovery of embryonic tissues, using a pipette (i.e. P200 for 2D-assays and P1000 for intact PSM), each sample was carefully loaded into the microfluidics device, which was already coated with fibronectin, immersed in a buffer of PBS and PenStrep, and degassed. Each sample inlet was plugged with a PDMS-filled PTFE tubing immediately after sample loading. Unused inlets, if any, were also plugged.

Flow rate of drug/medium was set to 20 uL/hr and the tubings were carefully inserted into the drug/medium inlets in the PDMS device while it is immersed in buffer. The tubings connected to the syringes with medium were inserted first, and the tubing connected to the syringe with the drug was inserted last. A 15-min timer was started after insertion of the drug tubing. PDMS device was then removed from the buffer and excess liquid on the cover glass was removed carefully with lint-free wipes (Kimberly-Clark). The PDMS device, with more than 1 m of attached tubings, was carefully placed inside a pre-equilibriated microfluidics holder (EMBL Mechanical Workshop, Figure S12) customized to fit the 70 mm x 70 mm cover glass (Marienfeld 0107999 098) and some PTFE tubing. Putting some of the tubing inside the microfluidics holder was necessary to equilibriate the drug/medium to desired environmental conditions before they are perfused into the microfluidics device. The cover glass was secured in place with grease and a U-shaped metal clamp. The microfluidics holder was then carefully mounted on the stage of the microscope, and the end of the outlet tubings were placed in a beaker.

Fifteen minutes after insertion of the drug tubing in the PDMS device, each pump was tilted on its side and was equilibriated for another 15 mins. Afterwards, the pump mounting the syringes with the drug and DMSO control was turned off, and the pump mounting the syringes with the medium was set to flow rate of 60 uL/hr. Samples were equilibrated at these conditions for at least 30 mins before start of imaging/entrainment.

### F. Setting up automated pumping

Entrainment via periodic pulses of drug was performed through alternate perfusion of medium and drug into the microfluidics device. Perfusion of drug/medium was done using syringes mounted on programmable syringe pumps (World Precision Instruments, AL-400). Diameter of syringe (14.5 mm for 10-mL syringe, BD Luer-Lok REF 300912) was accordingly set to match a defined flow rate and a specified volume of solution to be perfused with the duration of perfusion. Standard pumping programs of medium and drug/control are summarized in Tables S5 and S6, respectively, considering an entrainment experiment with *T*_*zeit*_ = 170 mins.

### G. Confocal microscopy

For most of the experiments here, samples were imaged in an LSM 780 laser-scanning microscope (Carl Zeiss Microscopy) fitted with an incubation chamber (EMBL Mechanical Workshop). A Plan-Apochromat 20x air objective with a numerical aperture (NA) of 0.8 (Carl Zeiss Microscopy) was used for imaging, and the zoom was set to 0.6. Three z-stacks (spacing: 8 um) were scanned for each sample every 10 mins to acquire timelapse movies. Imaging of multiple samples in multiple locations was done with a motorized stage, controlled using Zen Black software (Carl Zeiss Microscopy), and automated using a VBA macro developed by Antonio Politi (77), which is available at https://git.embl.de/grp-ellenberg/mypic. The dimension of the images was 512 pixels x 512 pixels, with a pixel size of 1.38 um and bit depth of 16-bit. Detection of drug pulses, using Cascade Blue (excited with 405 nm) added to the solution, was also done every 10 mins with lower image resolution: 1 z-stack, 32 pixel x 32 pixel (pixel size = 22.14 um). Imaging and automated pumping of drug/medium through the microfluidics device were started simultaneously.

**Fig. S12.**
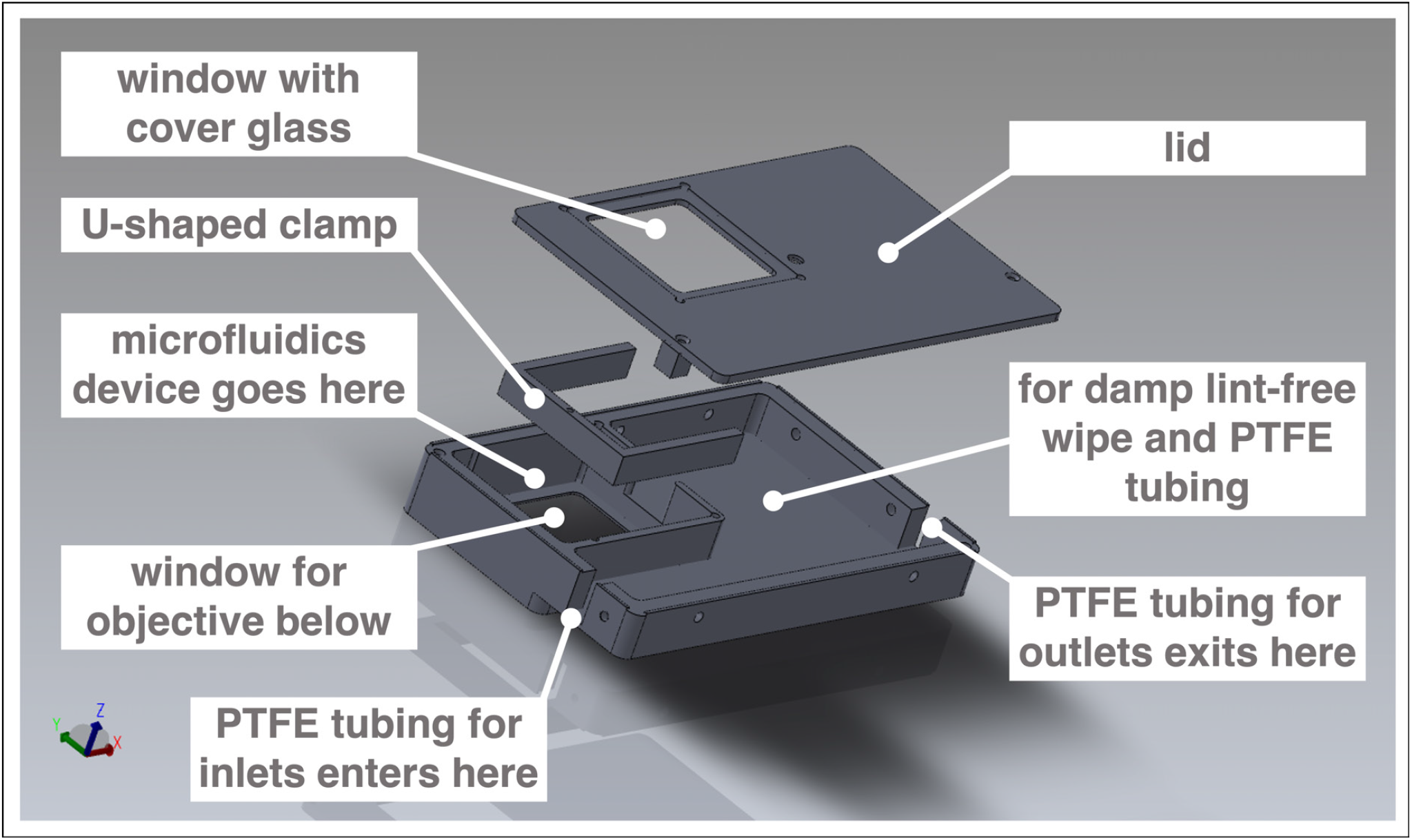
Customized box to mount microfluidics device on microscope for simultaneous culture, entrainment, and live imaging. Computer-aided design (CAD) of metal box customized to hold a microfluidics device bonded to a 70 mm x 70 mm cover glass. The box fits the stage of an LSM 780 laser-scanning microscope (Carl Zeiss Microscopy). Design by Katharina Sonnen, the EMBL Mechanical Design Office, and the EMBL Mechanical Workshop.

**Table S5.** Pumping program of medium for entrainment to 170-min periodic pulses of drug.

| Phase | Function | Rate | Volume | Remark |
| --- | --- | --- | --- | --- |
| PH:01 | LP:ST |  |  |  |
| PH:02 | RATE | 60 uL/hr | 140 uL | perfuse medium for 140 mins |
| PH:03 | LP:ST |  |  |  |
| PH:04 | PS:60 |  |  | pause for 60 secs |
| PH:05 | LP:30 |  |  | loop PH:04 30 times (pause for 30 mins = 60 secs × 30) |
| PH:06 | LP:30 |  |  | loop PH:02 to PH:05 30 times (total: 30 periodic pulses) |
| PH:07 | STOP |  |  |  |

**Table S6.** Pumping program of drug/control for entrainment to 170-min periodic pulses of drug.

| Phase | Function | Rate | Volume | Remark |
| --- | --- | --- | --- | --- |
| PH:01 | LP:ST |  |  |  |
| PH:02 | LP:ST |  |  |  |
| PH:03 | LP:ST |  |  |  |
| PH:04 | PS:60 |  |  | pause for 60 secs |
| PH:05 | LP:70 | | | loop PH:04 70 times (pause for 70 mins = 60 secs × 70) |
| PH:06 | LP:02 | | | loop PH:04 to PH:05 2x (pause 140 mins = 70 mins × 2) |
| PH:07 | RATE | 60 uL/hr | 30 uL | perfuse drug/control for 30 mins |
| PH:08 | LP:30 |  |  | loop PH:02 to PH:07 30 times (total: 30 periodic pulses) |
| PH:09 | STOP |  |  |  |

### H. Data analysis

Details on the coarse-graining strategy and analyses of entrainment dynamics are specified here.

#### Extracting timeseries from global intensity of 2D-assays for subsequent analyses

To extract the timeseries corresponding to the segmentation clock (Figure 2, S1A), global intensity analysis of timelapse fluorescence imaging was done using Fiji (78). Z-stacks were first projected based on maximum intensity [in Fiji: `Image Stacks > Z Project`]. Subsequently, the timeseries was obtained by plotting the z-axis profile [in Fiji: `Image > Stacks > Plot Z-axis Profile`]. Timeseries of replicate samples were compiled in a .txt file for subsequent analyses.

To extract timeseries corresponding to signaling oscillations localized at the center of 2D-assays, samples that were centered in the field of view were considered. A 50 pixel x 50 pixel oval region of interest (ROI) was specified at the center (x = 256 and y = 256) of a re-oriented (and registered, if necessary) 2D-assay timelapse file [in Fiji: `Edit > Selection Specify`]. Timeseries was extracted after specifying the center ROI [in Fiji: `Image > Stacks > Plot Z-axis Profile`]. Timeseries of replicate samples were compiled in a .txt file for subsequent analyses.

#### Definition of Kuramoto order parameter

We follow the definition from (36) : introducing the mean field variable

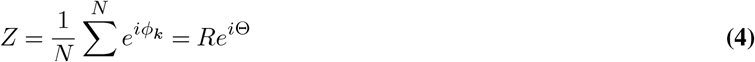

(see also (49), the first Kuramoto order parameter is defined by

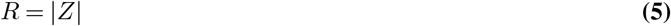

#### Monitoring period-locking and phase-locking

Entrainment was evaluated based on period-locking and phase-locking of the signaling oscillations to the periodic drug pulses. Oscillatory components were extracted from timeseries using a wavelet analysis workflow that was developed by Gregor Mönke, which was recently implemented as a Python-based standalone software (79) available at https://github.com/tensionhead/pyBOAT. In this workflow, timeseries was first detrended using a sinc filter and then subjected to continuous wavelet transform. Time-resolved frequency analysis was done by cross-correlating the signal to wavelet functions of known frequencies, generating a power spectrum. A high power score was assigned to wavelets that correlated well with the signal relative to white noise. Instantaneous period and phase were extracted upon evaluation of the power spectrum along a ridge tracing wavelet with maximum power for every timepoint.

Phase dynamics of signaling oscillations, upon subjecting them to periodic perturbation, were analyzed using stroboscopic maps (38, 39, 53). Briefly, the phase difference (Δ *φ*) was defined as:

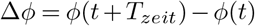

where *t* is the time of perturbation and *T*_*zeit*_ is the *zeitgeber* period (i.e. one cycle after time = *t*). *φ*(*t*) and *φ*(*t* + *T*_*zeit*_) denote the `old_phases` and their corresponding `new_phases`, respectively. The stroboscopic maps were then plotted as `new_phases` versus `old_phases` (for scheme, please refer to Figure 6A). The centroid, marking the entrainment phase (*φ*_*ent*_), was determined considering only phase-locked samples (where the difference between a sample’s phase at the time of the final drug pulse considered and i ts phase one drug pulse before is less than *π /*8). This was quantified from the average phases of the final (`old_phase`,`new_phase`) pairs considered, and the circular standard deviation (circSD) was calculated using the formula:

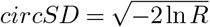

where *R* is the first Kuramoto order parameter. As wavelets only partially overlap the signal at the edges of the timeseries, resulting in deviations from true phase values (79), the first and last pulse pairs were not considered in the generation of stroboscopic maps. Polar plots were also generated summarizing the instantaneous phase of replicate samples and their first Kuramoto order parameter as shown in Figure 4D, Figure 5C, and Figure S5B. The Python code is available as a Jupyter notebook (.ipynb) at https://github.com/PGLSanchez/EMBL_OscillationsAnalysis/tree/master/EntrainmentAnalysis. This code uses Matplotlib (80), NumPy (81, 82), pandas (83), scikit-image (84), SciPy (85), and seaborn (86).

Detrended timeseries of replicate samples were in some part of the study represented as a heatmap using PlotTwist (50), as shown in Figure 4A. Average periods (from 650 mins to 850 mins after start of experiment) were meanwhile plotted using PlotsOfData (87), as shown in Figures 3C-D and Figure S3. These apps are available at https://huygens.science.uva.nl/PlotTwist/ and at https://huygens.science.uva.nl/PlotsOfData/, respectively. To calculate p-values, two-tailed test for absolute difference between medians was done via a randomization method using PlotsOfDifferences (47). While not recommended, p-values were also calculated similarly for conditions with n < 10. PlotsOfDifferences is available at https://huygens.science.uva.nl/PlotsOfDifferences/.

#### Generating wavelet movies

Period and phase wavelet movies were generated using the Wavelet Processing and Export workflow developed by Gregor Mönke, which runs on the EMBL cluster and is implemented in Galaxy with technical assistance from Jelle Scholtalbers (EMBL Genome Biology Computational Support). This workflow extracts timeseries of every pixel in a timelapse movie and subjects them to sinc filter-based detrending and subsequent continuous wavelet transform (79). This results in extraction of instantaneous period and phase of each pixel, recovering period and phase wavelet movies corresponding to the input timelapse movie (for scheme, please refer to Figure S11A). The workflow is available at https://github.com/tensionhead/SpyBOAT and can be used via a public Galaxy server at https://usegalaxy.eu/root?tool_id=/spyboat. Settings used to generate wavelet movies in this study were: sigma of 8.0, sample interval of 10 mins, period range from 100 to 250 mins, and number of periods analyzed of 151.

#### Examining period gradient in 2D-assays

Re-oriented 2D-assays (dorsal side up) and their corresponding period and phase wavelet movies were used for the analyses. Temporal evolution of the period gradient during the course of entrainment experiments was evaluated from the period wavelet movies of the 2D-assays. As the period wavelet movies also contained wavelet transformations for pixels in the background, a binary mask was first created to differentiate pixels corresponding to signal. To create the mask, re-oriented (and registered, if necessary) timelapse movies of 2D-assays were blurred using a Gaussian blur [in Fiji: `Process > Filters Gaussian Blur` (sigma radius: 8, scaled units in microns)]. Then, signal was specified by thresholding [in Fiji: `Image Adjust > Threshold` (Default method, dark bakground)]. After thresholding, pixels corresponding to the signal were assigned a value of 255, while those corresponding to the background were assigned a value of 0. If opposite, the values were inverted [in Fiji: `Edit > Invert`]. Then, the binary mask (signal = 1 and background = 0) was created by dividing all values by 255 [in Fiji: `Process > Math > Divide` (value: 255)], and was used to mask the period wavelet movie.

**Fig. S13.**
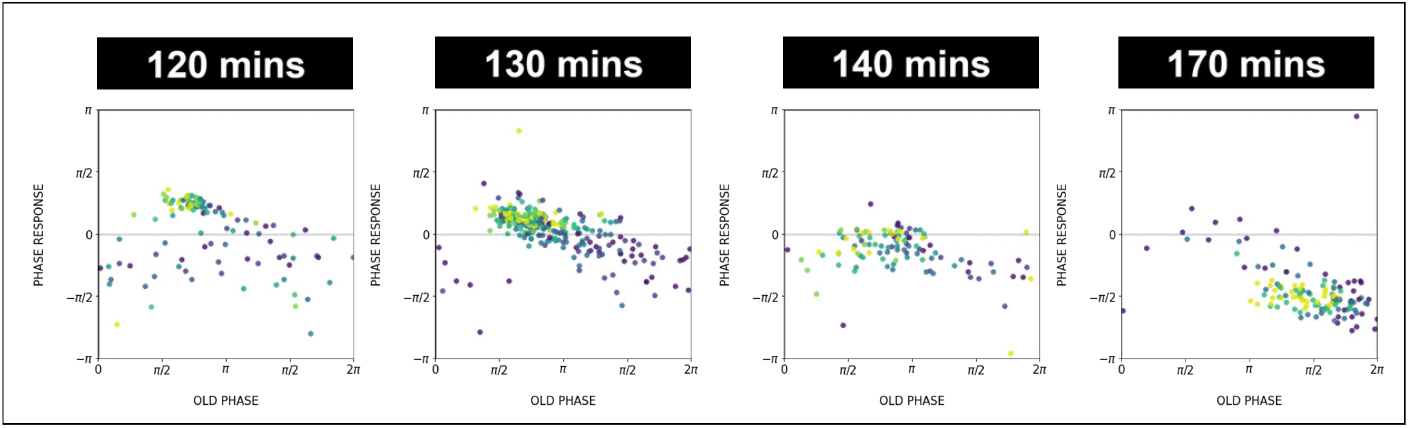
Phase response as a function of time. We use the color code for pulse number similar to Fig. 6 (from purple for early pulses to yellow for later pulses). In the PRC from the 120 min (170 min) experiment later data points are located higher (lower) than earlier data points, consistent with the change of the intrinsic period of oscillations during entrainment. The adjustment of the intrinsic period decreases the detuning, causing the vertical shift of the phase response curve.

**Fig. S14.**
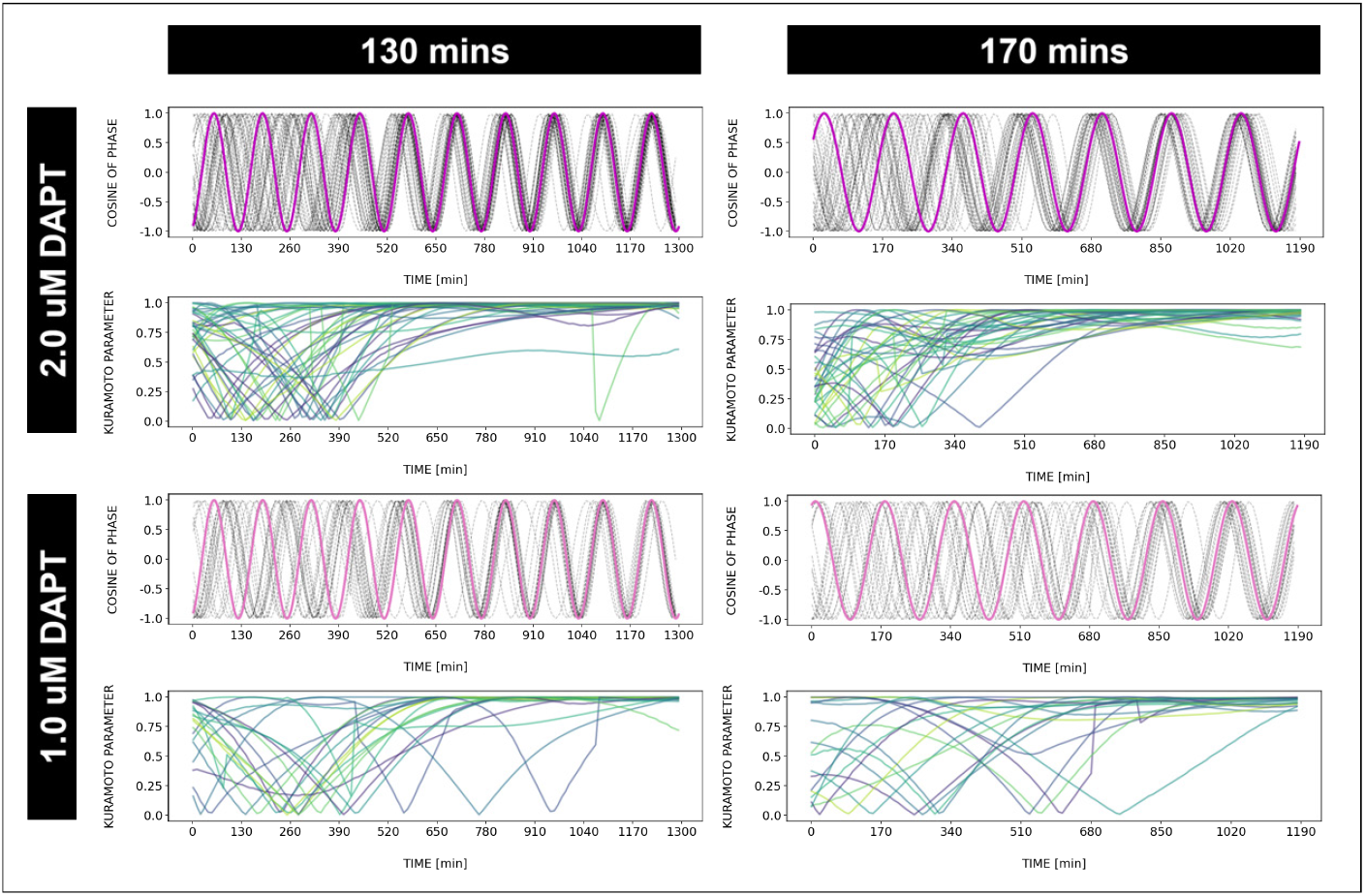
Dynamics and variability of entrainment. We represent the *zeitgeber* signal as a continuous uniformly increasing phase (*‘zeitgeber time’*) with period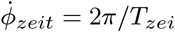. The initial condition for *φ*_*zeit*_ is chosen so that the *zeitgeber* phase at the moment of the last pulse is matching the experimental entrainment phase for each *T*_*zeit*_. We plot the corresponding cos *φ*(*t*) for each sample (dotted lines) and the zeitgeber phase (solid lines). To quantify how well each sample is following the *zeitgeber*, we compute the Kuramoto parameter: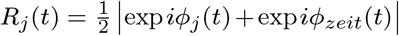, where *φ*_*j*_ (*t*) is the phase of sample *j*. Convergence to 1 of the Kuramoto parameter *R*_*j*_ indicates the establishment of a stable phase relation between the sample and *zeitgeber* pulses, indicative of entrainment. As seen from the plots, at lower concentration of DAPT samples take a longer time on average to reach entrainment.

## Supplementary Note 2: Theoretical methods

### A. PRC correction and computation

PRCs at different entrainment periods appear shifted with respect to one another. To correct for this, we fitted the PRCs at entrainment periods *T*_*zeit*_ = (120, 130, 140, 170) mins (those for which we have enough experimental data points to build a continuous curve) using 3 Fourier modes (Figure 8A1) and manually chose the phase shift optimizing their overlap for Figure 8A2. The shift values used were (*−* 0.45, *−* 0.25, 0, 0.45) for *T*_*zeit*_ = (120, 130, 140, 170) mins respectively. The data points shown in Figure 8B are used for Monte Carlo optimization (see next section). Our choice was later validated by recomputation of PRCs and comparison with data in Figure 8G. We also show corresponding PTCs in Figure S17.

### B. Model

Our model (Figure S15A) is based on simple modifications of the classical Poincaré oscillator, otherwise called Radial Isochron Cycle model (RIC). The main motivation is to find the simplest possible model able to reproduce the general shape of the experimental PRC, with both a flat and a negative region, while being analytical with simple isochrons. The full description and analytical calculations for this model will be described elsewhere, here we outline the basic equations and some general properties of the model. For PRC computations, we will assume that a perturbation is a horizontal shift of magnitude *A* (Figure S15B).

To flatten the PRC for half the cycle, we first modify the limit cycle of the standard RIC model into an ellipse, keeping the origin as one of the foci. This is done through the introduction of a parameter *λ*, so that the corresponding equation of the ellipse is

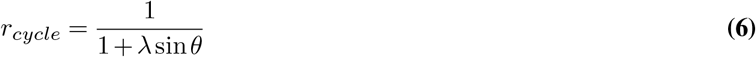

combined with radial equidistant isochrons, 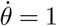, where *θ* is the polar angle. This is however not enough to define the entire flow in the plane since one needs to specify how *r* converges on this cycle. We thus impose a radially uniform convergence rate equal to 1 for the radius, which leads to the following differential equation in polar coordinates:

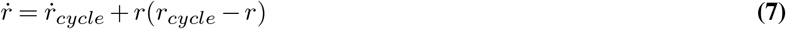

where 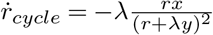 (again assuming 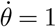). This leads to the following equations in cartesian coordinates:

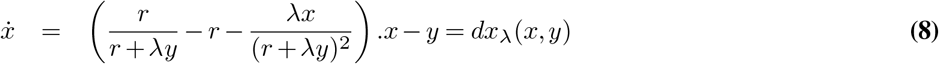

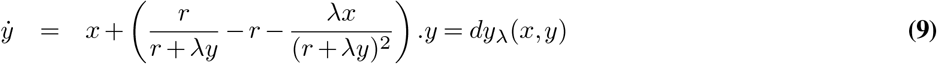

We name this model Elliptic Radial Isochron Cycle, or ERIC. An important feature of ERIC is that the angle in the plane is the phase of the oscillator, in particular since the phase is defined by the planar angle, the PRC following a horizontal perturbation of size *A* towards the right can be computed in a straightforward way and is :

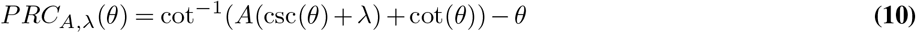

As said above, the effect of introducing *λ* is to smoothly flatten the PRC over half of the cycle, as illustrated in Figure S16A. We then introduce a second modification, allowing us to rescale the portion of the cycle where the PRC is flattened. We modify the equations by introducing a “speeding factor” *s* so that

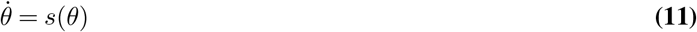

This will keep isochrons radial but will change their spacing. With the elliptic limit cycle, this leads to the differential equation in cartesian coordinates

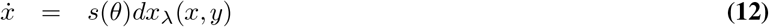

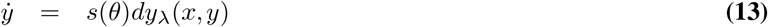

We name this class of model Elliptic Radial Isochron Cycle with Acceleration, or ERICA. For simplicity, and to keep the system analytical, we restricted ourselves first to *s* functions linear by piece, i.e. *s* = *s*_*\**_ *>* 1 for one sector and *s* = 1 otherwise. The sped up sector is centered at angle *α* and has width *β*, so that the modified period of the cycle is *T*_*s*_*\** = *β/s*_*\**_ + (2*π − β*). It is also convenient to define the rescaled angular velocity 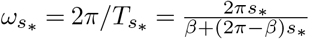. From there, the phase of the cycle as a function of the angle *θ* in the plane is the simple linear transformation (defining *θ*_0_ = *α − β/*2) :

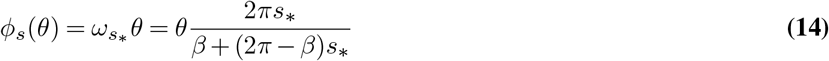

for 0 *< θ < θ*_0_,

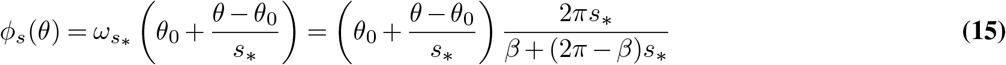

for *θ*_0_ *< θ < θ*_0_ + *β* and

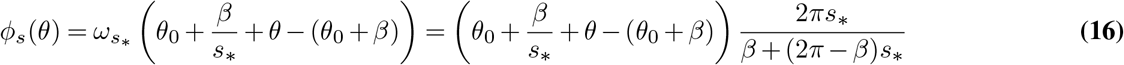

for *θ*_0_ + *β < θ <* 2*π*. Those functions ensure that the rate of phase evolution in sector *θ*_0_ *< θ < θ*_0_ + *β* is 1*/s* times the rate in the other sectors (compare *θ* coefficients in Eq. 14-16), that angle 0 in *θ* is phase *φ*_*s*_ = 0, and that phase is continuous so that 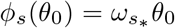 and 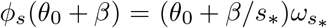. Notice also that for *θ* = 2*π* we get from Eq.16 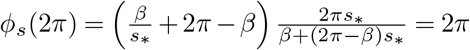 as expected after one full cycle.

A full study of the possible behaviours of the ERICA model will be published elsewhere. Figure S16 illustrates different shapes of PRC obtained by varying *λ, s*_*\**_, *α*, and *β* independently, for various choices of sped up sectors.

For optimization purpose, it is easier to numerically compute the PRC in the following way. We consider an ensemble of angles *θ*_*i*_ linearly spaced on the interval [0, 2*π*]. We then compute the corresponding position on the ERIC limit cycle, and compute numerically the angle 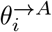 of the point at distance *A* on the right. By definition of ERICA, the PRC for each index *i* then is

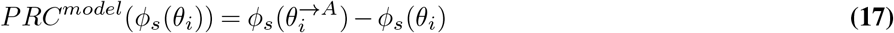

We checked that this PRC coincides both with the one computed from the integration of the ODEs (simulating the full stroboscopic map procedure) and with the analytical expression. To fit the experimental PRC, we run Monte Carlo simulations to optimize parameters *A, λ, s*, sped up sector location *α* and width *β*, and the location of the zero-phase reference point on the cycle *φ*_0_. We minimize *χ*^2^:

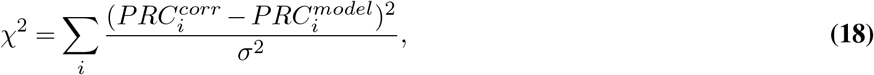

where 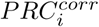 are data points of the corrected experimental PRC and *σ* is the noise, estimated as the root mean square error of a Fourier fit to data (see Figure 8A). We use Markov chains to generate distributions of parameters and take the average values to plot the model limit cycle and PRC. The optimized parameter values are: *A* = 0.43, *λ* = 0.53, *s* = 5.64, *α* = 1.51*π, β* = 1.15*π, φ*_0_ = 1.17*π*.

### C. Evidence for intrinsic period changes

The PRC shifts are interpreted as change of period of the intrinsic oscillator because of entrainment.

We first checked that such PRC shifts are really needed to explain data. To do this, we computed an average PRC by only taking into account entrainment periods between 130 and 150 mins. If we assume there is no period change, we can compute the maximum entrainment period compatible with such PRC, and found it is equal to 170 mins. However it is clear from our data that we can go beyond this entrainment period, confirming the shift of PRC, in agreement with a change of intrinsic period. We further have two evidences for such changes from release experiments. In those experiments, we entrain the oscillators for 3 cycles at 170 mins, then let it go, and measure both the phase and instantaneous periods of the oscillator (Figure S8). When we measure the period, we first find it takes several cycles to come back to the intrinsic period, showing a long lived effect, unlikely coming from some artefact of period computation (Figure S8). A more direct way to estimate period change is to compute the return maps after the last pulse of DAPT. In Figure S17A we consider all release experiments, which were done with *zeitgeber* period of 170 mins. We take the phase one full cycle after the last pulse, and compute the return map from this phase (so for the first full cycle with no pulses). From this, we can compute the length of the cycle to have an average phase difference of 0 (corresponding to the period two full cycles after the last pulse). We found a cycle length of 150 minutes, compatible with the instantaneous period measurement and incompatible with a fast return to a 140 minutes intrinsic period.

We further show the Arnold tongue computed numerically with a similar PRC (1 : 1 entrainment region) and corresponding isophases assuming a fixed intrinsic period of 140 minutes in Figure S17B. We see much narrower width clearly incompatible with the data, again confirming the need for an intrinsic period change.

The last evidence is both more indirect but more mathematically grounded. It comes from the sigmoidal shape of the experimental entrainment phase as a function of *zeitgeber* period (Figure 7B, 8E). In a nutshell, flatness of this curve is incompatible with the classical PRC theory, but can be easily explained if the internal period *T*_*osc*_ changes linearly with *T*_*zeit*_.

To see this, we start with the classical condition for stability of the entrainment phase (see e.g. (52)),

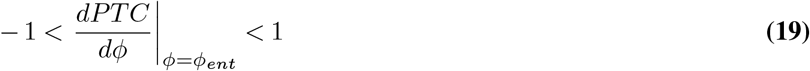

or equivalently :

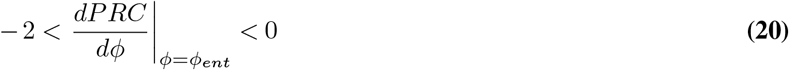

Now we also have by definition of the entrainment phase *φ*_*ent*_ :

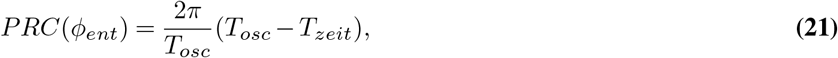

where the factor 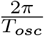 is the unit conversion from time to phase. This equation defines in an implicit way a function *φ*_*ent*_(*T*_*zeit*_), and the corresponding experimental curve is plotted in Figure 8E for our data. Taking the derivative on both sides with respect to the *zeitgeber* period, we get :

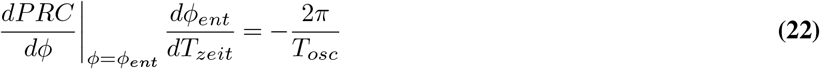

This allows to implicitly define the curve *φ*_*ent*_(*T*_*zeit*_) with the help of the *PRC*:

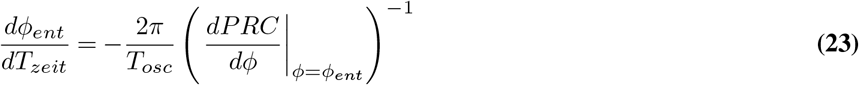

There is a geometric interpretation here : the curve *φ*_*ent*_(*T*_*zeit*_) is in fact a (rescaled) +*π/*2 rotation of the PRC! This can be seen here because Eq. 23 combines a the mirror image of the PRC along the *y* axis (minus sign) with a mirror image along the first diagonal (power 1 which corresponds to the inversion function exchanging *φ* and *PRC* (*φ, PRC*(*φ*))→ (*PRC*(*φ*), *φ*)). We know from classical group theory that two mirror images give one rotation with an angle equal to twice the angle between the axis (so here 2*π/*4 = *− π/*2). For instance, if the PRC is sinusoidal, *φ*_*ent*_(*T*_*zeit*_) looks itself like (half a period) of a vertical sinusoidal, as can be clearly seen e.g. in (54). In Figure 8C, we further illustrate this rotation using the PRC computed from the data: the PRC rotated by *π/*2 is compared with the curve *φ*_*ent*_(*T*_*zeit*_) computed numerically from it, showing perfect agreement.

Now, since the geometry of the PRC is constrained by Eq. 20, this in turn imposes geometric constraints on *φ*_*ent*_(*T*_*zeit*_), e.g. combining Eqs. 20-23 and counting time in units of the intrinsic period *T*_*osc*_, 2*π/T*_*osc*_ = 1, we get

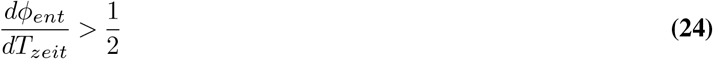

This defines an absolute, *minimum* slope of the curve *φ*_*ent*_(*T*_*zeit*_). However, we see experimentally in Figure 8E that the entrainment phase is becoming almost constant for low and high values of the entrainment period, indicating a zero slope, which thus is theoretically impossible from Eq. 24.

Now the simplest way to reconcile this observation with this calculation is to assume that the intrinsic period *T*_*osc*_ in fact depends on the *zeitgeber*. We then get a generalized version of Eq. 22 with changing intrinsic period

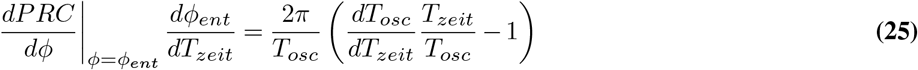

giving the new implicit definition

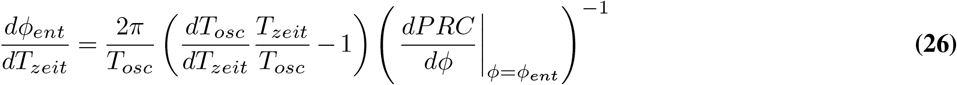

Now we see that when 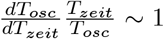, the right hand side of Eq. 26 can be very small, so that one can get 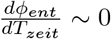, meaning that the entrainment phase barely changes as observed experimentally in Figure 8E. This effect will also come with a considerable enlargement of the Arnold tongue as discussed below. Also we notice that 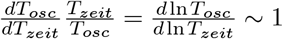 could be indicative of a mechanism where the oscillator adapts its intrinsic period (here to the *zeitgeber* period).

Practically, we obtain the *T*_*osc*_(*T*_*zeit*_) dependency by fitting the stroboscopic maps and entrainment phase from all experiments with 2.0 uM DAPT, using the PTC of our optimized model (Figure S17D-E). The values of *T*_*osc*_ are shown as data points in Figures 8D and S17F.

### D. Arnold tongue computations

To build the Arnold tongues, we first need to interpolate the period changes for period values not used in entrainment experiments. We used cubic spline interpolation to draw Figure 8D. For periods outside of the range of entrainment, in the absence of data some arbitrary choices have to be made. We know experimentally that entrainment does not occur below *∼* 120 mins and above *∼* 200mins which indicates that 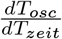 is becoming smaller around those *zeitgeber* periods. More biologically, this indicates that the system does not adjust to any entrainment period. Again some arbitrary choices have to be made, but based on those constraints and our own interpolation, a cut-off for *T*_*osc*_ of *±*20% of the natural period of 140 mins was assumed, and is further consistent with the relative period changes observed experimentally in zebrafish segmentation clock mutants (21) and theoretically derived from delayed coupled models (88). With such cut-off, the obtained entrainment range of *zeitgeber* periods is from 118 to 200 mins.

Computation of Arnold tongue and isophases in Figure 8F was done by iterations of the return map Eq. 2 and detection of fixed points, using as the intrinsic period the interpolation and extrapolation showed in Figure S17F.

Lastly, we can use this model to compute entrainment phase for different DAPT concentrations. For the main concentration of 2.0 uM DAPT, we have the optimized amplitude of perturbation *A* = 0.43. Assuming that the DAPT concentration is reflected in the strength of perturbation, we can find cross-sections of the isophases at different values of *A* to have excellent agreement with experiments, as illustrated in Figure S17G. We take *A* = 0.55 for 3.0 uM, *A* = 0.31 for 1.0 uM, and *A* = 0.13 for 0.5 uM DAPT, as shown on the Arnold tongue in Figure 8F. We capture both the presence/absence of entrainment and the *φ*_*ent*_(*T*_*zeit*_) relation. In particular, we see that for lower DAPT concentrations, the range of entrainment is smaller but we still get plateaus in entrainment phase consistent with the change of period we computed.

**Fig. S15.**
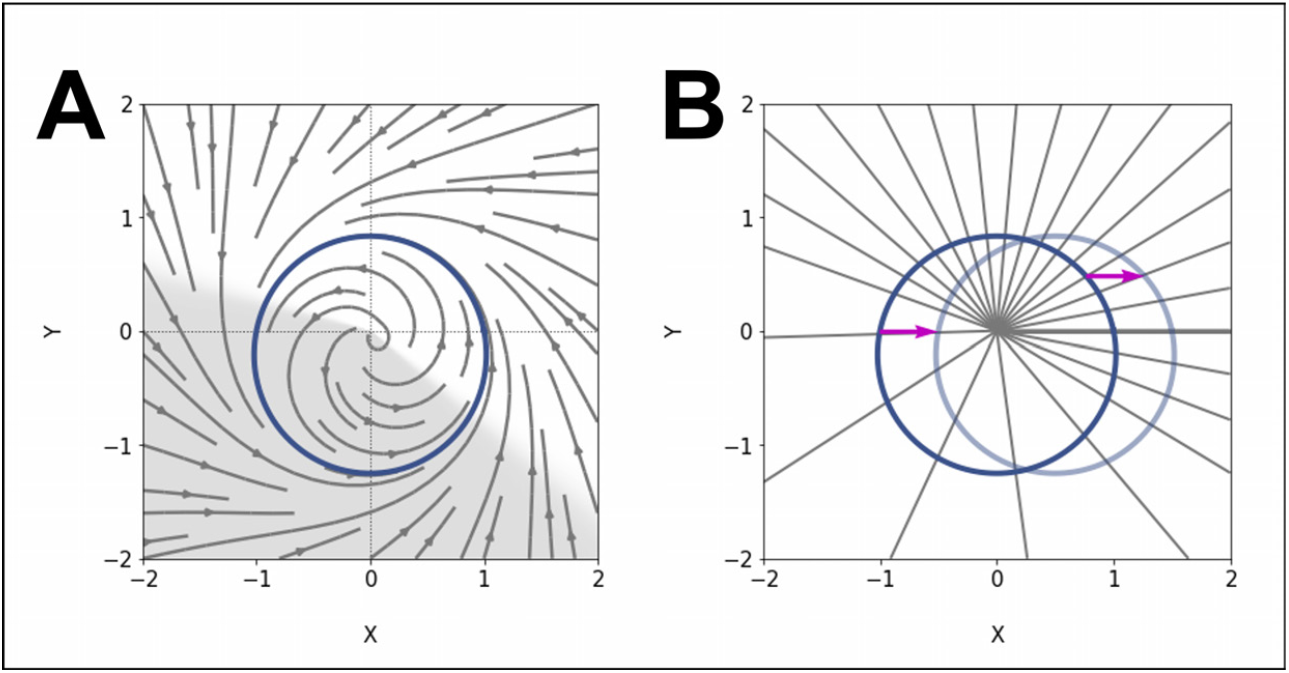
ERICA model. (**A**) Our model consists of an elliptic limit cycle (blue) with increased angular velocity in one sector (shaded region). (**B**) The effect of such acceleration is to change the spacing of isochrons (radial lines). We compute the PRC of the model by introducing a perturbation (arrow) at points on the limit cycle and looking at the starting and ending isochrons.

**Fig. S16.**
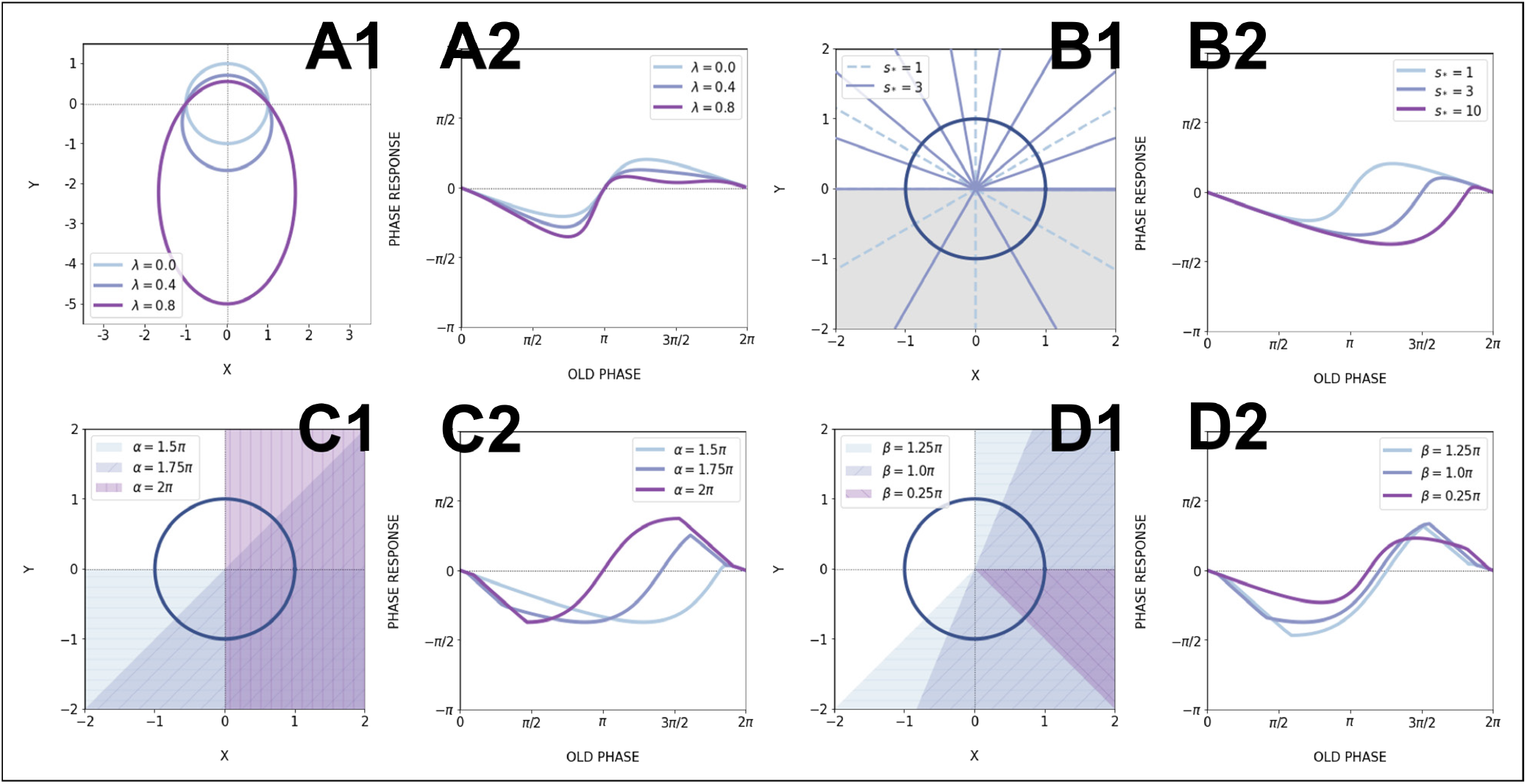
Effects of model parameters on the PRC. (**A**) Effect of eccentricity *λ*: (**A1**) Increasing *λ* makes the limit cycle more elongated, while keeping the upper focus at the origin. (**A2**) The positive part of the PRC is then flattened; for all PRCs here *s*_*\**_ = 1 (no acceleration). (**B**) Effect of speeding up *s*_*\**_ : (**B1**) When *s*_*\**_ is increased, the spacing between isochrons in the sped up region increases, so this region contains less and less isochrons. Shown here is the case when the acceleration happens in the lower half-plane (*α* = 1.5*π, β* = *π*), which is shaded in grey. (**B2**) The effect on the PRC is to shrink the portion where the oscillator is sped up, thus emphasizing the other part of the curve. Here *λ* = 0. (**C, D**) Effects of the sped up sector parameters *α, β*: the location and width of the sector determine which parts of the PRC get rescaled. (**C1**) The shaded regions are sectors with different values of *α*, and *β* = *π*. (**C2**) The corresponding PRCs with *λ* = 0, *s*_*\**_ = 10. (**D1**) Sped up sectors located at *α* = 15*/*8*π* with different widths *β*. (**D2**) The corresponding PRCs for the case *λ* = 0, *s*_*\**_ = 10. All PRCs in this figure were computed with perturbation amplitude *A* = 0.6.

**Fig. S17.**
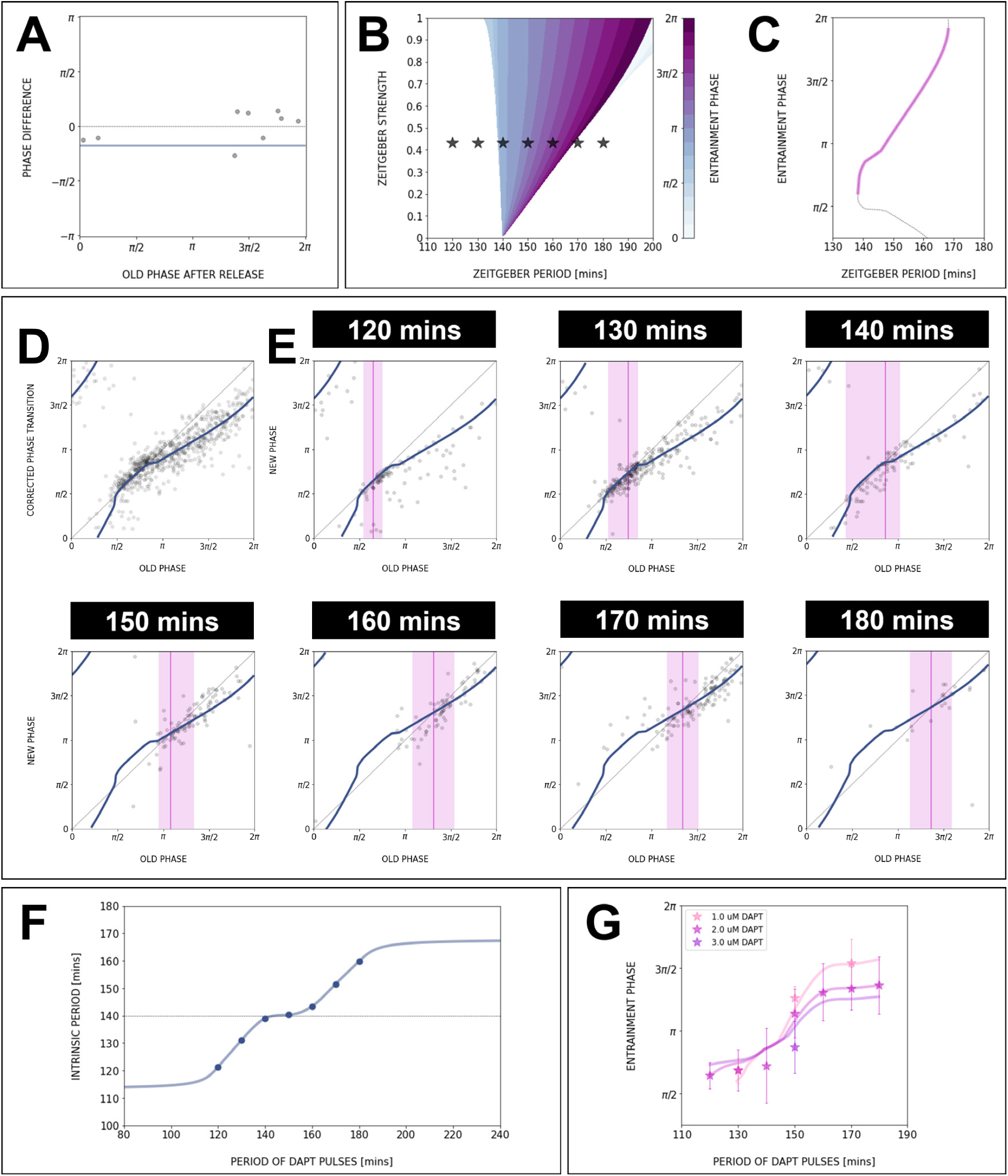
Data for the PRC/PTC, stroboscopic maps and entrainment phase can be fully captured with our model and changing intrinsic period. (**A**) Stroboscopic map data from release experiments allows to estimate the intrinsic oscillation period during entrainment. With *T*_*osc*_ = 150 mins, the phase difference between the end and the beginning of the first full cycle with no perturbations is on average 0 (2*π*). The data points are from different release experiments with *T*_*zeit*_ = 170 mins. If the natural period of 140 mins is used, the average phase difference is 0.17*π* (solid line). (**B, C**) Constant intrinsic period *T*_*osc*_ is not consistent with experimental data: (**B**) The 1 : 1 Arnold tongue, computed using our model and *T*_*osc*_ = 140 mins, is much narrower than the experimental entrainment range. Stars correspond to experimental conditions (*T*_*osc*_, *A*) where entrainment was observed. (**C**) Entrainment phase *φ*_*ent*_(*T*_*zeit*_) numerically computed with the optimized PRC from Figure 8C, assuming constant *T*_*osc*_ = 140 min (which also corresponds to a cross-section of the isophases in **B**). Clearly, the slope of the curve is much higher than in the data. For comparison, we also plot the Figure 8C PRC rotated by *π/*2, showing perfect overlap. (**D**) PTC of the optimized model (equivalent to the PRC shown in Figure 8C). (**E**) Using the PTC from **D**, we fit the stroboscopic maps data for all periods *T*_*zeit*_ by choosing the *T*_*osc*_(*T*_*zeit*_) that gives the right detuning. The narrow magenta lines indicate the entrainment phase, the fixed point of each fitted stroboscopic map. The shaded magenta regions show the experimental range for entrainment phase. (**F**) Extrapolation of the intrinsic oscillator period *T*_*osc*_ as a function of zeitgeber period *T*_*zeit*_. (**G**) Cross-sections of the isophases in Figure 8F, calculated with the model and the extrapolated curve for *T*_*osc*_(*T*_*zeit*_), give excellent agreement with the entrainment phase data for different concentrations of DAPT.

## Supplementary Note 3: Movies

- E10.5 2D-assay, expressing LuVeLu, subjected to 130-min periodic pulses of DMSO control, with corresponding period and phase wavelet movies, available at https://github.com/PGLSanchez/EMBL-files/blob/master/MOVIES/SO_2.0D_130mins_CTRL.avi
- E10.5 2D-assay, expressing LuVeLu, subjected to 130-min periodic pulses of 2 uM DAPT, with corresponding period and phase wavelet movies, available at https://github.com/PGLSanchez/EMBL-files/blob/master/MOVIES/SO_2.0D_130mins_DAPT.avi
- E10.5 2D-assay, expressing LuVeLu, subjected to 170-min periodic pulses of DMSO control, with corresponding period and phase wavelet movies, available at https://github.com/PGLSanchez/EMBL-files/blob/master/MOVIES/SO_2.0D_170mins_CTRL.avi
- E10.5 2D-assay, expressing LuVeLu, subjected to 170-min periodic pulses of 2 uM DAPT, with corresponding period and phase wavelet movies, available at https://github.com/PGLSanchez/EMBL-files/blob/master/MOVIES/SO_2.0D_170mins_DAPT.avi
- E10.5 2D-assay, expressing LuVeLu, subjected to 170-min periodic pulses of 2 uM DAPT (or DMSO for control), with corresponding timeseries obtained using a global ROI spanning the entire field of view, available at https://youtu.be/fRHsHYU_H2Q
- E10.5 2D-assay, expressing Axin2-linker-Achilles, subjected to 170-min periodic pulses of 2 uM DAPT (or DMSO for control) available at https://youtu.be/edFczx_-9hM
- E10.5 2D-assay, expressing Mesp2-GFP, subjected to 170-min periodic pulses of 2 uM DAPT (or DMSO for control) available at https://youtu.be/tQeBk0_U_Qo
- Period gradient of LuVeLu in E10.5 2D-assay subjected to 170-min periodic pulses of 2 uM DAPT (or DMSO for control) available at https://youtu.be/3Y53OhXacKI

## Supplementary Note 4: Text files

- Text (.txt) files containing timeseries from microfluidics-based entrainment experiments, available at https://github.com/PGLSanchez/EMBL-files/tree/master/ENTRAINMENT-timeseries

